# TractEM: Fast Protocols for Whole Brain Deterministic Tractography-Based White Matter Atlas

**DOI:** 10.1101/651935

**Authors:** Roza G. Bayrak, Xuan Wang, Kurt G. Schilling, Jasmine M. Greer, Colin B. Hansen, Justin A. Blaber, Owen Williams, Lori L. Beason-Held, Susan M. Resnick, Baxter P. Rogers, Bennett A. Landman

**Author notes:** Corresponding author Email address (Roza G. Bayrak).

## Abstract

Reproducible identification of white matter tracts across subjects is essential for the study of structural connectivity of the human brain. The key challenges are anatomical differences between subjects and human rater subjectivity in labeling. Labeling white matter regions of interest presents many challenges due to the need to integrate both local and global information. Clearly communicating the human/manual processes to capture this information is cumbersome, yet essential to lay a solid foundation for comprehensive atlases. The state-of-the-art for white matter atlas is the single population-averaged Johns Hopkins Eve atlas. A critical bottleneck with the Eve atlas framework is that manual labeling time is extensive and peripheral white matter regions are conservatively labeled. In this work, we developed protocols that will facilitate manual virtual dissection of white matter pathways, with the goals to be anatomically accurate, intuitive, reproducible, and act as an initial stage to build an amenable knowledge base of neuroanatomical regions. We analyzed reproducibility of the fiber bundles and variability of human raters using DICE correlation coefficient, intraclass correlation coefficient, and root mean squared error. The protocols at their initial stage have shown promising results on both typical 3T research acquisition Baltimore Longitudinal Study of Aging and high-acquisition quality Human Connectome Project datasets. The TractEM manual labeling protocols allow for reconstruction of reproducible subject-specific fiber bundles across the brain. The protocols and sample results have been made available in open source to improve generalizability and reliability in collaboration.

## 1. Introduction

Human brain atlases have had a transformative role in the study of neuroscience. Evolving neuroimaging, image processing, and analysis techniques are framing the way that we understand brain anatomy. Modern brain atlases (Mazziotta et al., 2001; Hawrylycz et al., 2012; Amunts et al., 2013) combine population atlases with multi-modal structural and functional information. These atlases provide a detailed representation of the anatomy and are a fundamental component of the field of image analysis. Although most commonly used modern approaches tend to treat white matter as essentially homogeneous, most current atlases are incomplete without detailed white matter information. A whole-brain white matter (WM) atlas, for subject-specific population atlases, labeled with equivalent methodologies, tools and protocols is therefore a fundamental resource for integrative and comprehensive modern brain atlases.

The effort to create atlases is not new. The importance of a white matter atlas arises from the information that can be inferred from them. White matter connectivity stores substantial information about functional neuroanatomy (Sporns et al., 2005; Skudlarski et al., 2008; van den Heuvel et al., 2008; Greicius et al., 2009), longitudinal changes (Resnick et al., 2003), brain morphology (Lawes et al., 2008), abnormalities and their cognitive correlation (Gunning-Dixon and Raz, 2000), critical periods for neural development (Anderson et al., 2011), recovery and plasticity (Jiang et al., 2006; Dayan and Cohen, 2011), and cognitive functioning (Llinás et al., 1998). Nonetheless, existing white matter atlases are limited to population-averaged single atlases with extremely time consuming manual delineation (Mori et al., 2008), atlases lacking comprehensive coordinate information about white matter structures (960), atlases targeting selective tracts instead of the whole-brain (Wakana et al., 2004; Oishi et al., 2008; Adluru et al., 2016; Besseling et al., 2012; Garyfallidis et al., 2018; Nath et al., 2018; Zhang and Arfanakis, 2018) or atlases created with automated methods that rely on manual delineations (Garyfallidis et al., 2018; Wasserthal et al., 2018; Yeh et al., 2018).

While there is expert evaluated white matter atlases (Essayed et al., 2017; Yeh et al., 2018), they lack protocols detailing how to delineate white matter structures. The current state-of-the-art white matter atlas is the population-based single-subject Johns Hopkins University (JHU) atlas labeled using the Eve protocol (Mori et al., 2008; Oishi et al., 2008), using both diffusion tensor data and a T1-structural image of the brain. The JHU Eve atlas is based on diffusion weighted magnetic resonance imaging and subsequent diffusion tractography. Diffusion magnetic resonance imaging allows diffusion of water molecules to be visualized by sensitizing images in the direction of the field gradients. Diffusion tensor imaging uses this information to calculate anisotropy and therefore orientation information of the neural pathways. There are several limitations to the use of the Eve-based frameworks: (1) it is a population-averaged single atlas, (2) peripheral white matter regions are conservatively labeled and (3) averaging over many people causes blurring of small pathways. Additionally, (4) the atlas was generated using diffusion tensor imaging, which has been shown to have limitations in crossing fiber and complex regions (Oouchi et al., 2007), whereas a number of more advanced techniques successfully can overcome these limitations (Tuch, 2004; Descoteaux et al., 2007; Yeh et al., 2010).

In this work, we rectify the lack of manual labeling protocols that are robust to human subjectivity. This was achieved by creating protocols that are driven by strong anatomical prior. This strong prior is the state-of-the-art Eve white matter atlas that is obtained in consensus with many experts with neuroanatomical knowledge. The TractEM project aimed to allow expert/non-expert users to create and curate reproducible pathways across the brain. We developed TractEM and implemented a pipeline that allows tracking for 35 fiber tracts resulting in total 61 unique pathways (including L/R) in less than 6 hours with a methodology that can be applied generally to a large number of subjects. The protocols are tested on multiple datasets and are assessed for reproducibility. To the best of our knowledge, this is the first study that assesses human variability across minor and major pathways.

This method will serve as a valuable tool for the neuroimaging community. Additionally, the TractEM project in its initial stage curated a dataset that includes real human acquisitions, along with careful manual selection and definitions of ground truth streamlines. Subject-specific TractEM tractography data from the Human Connectome Project dataset is open to public use. We provided not only white matter volumes (or labels), but also streamlines themselves, a feature not provided by other open sourced datasets (Wasserthal et al., 2018; Zhang et al., 2018). Thus, TractEM will show value in region identification, normalization, anatomical validation, and even development of machine learning based tracking strategies (Poulin et al., 2019).

This paper is organized as follows. Section 2 describes the pipeline and methodology for the construction of tracts. Section 3 presents the results of intra-subject interrater tract reproducibility, tract variability, and rater reproducibility. Section 4 is reserved for general specific discussion, while section 5 offers a conclusion.

## 2. Materials and Methods

### 2.1. Data and Acquisition

The protocol was developed on datasets obtained from the Baltimore Longitudinal Study of Aging (BLSA) (Ferrucci, 2008) and Human Connectome Project (HCP) (Van Essen et al., 2012). The pipeline is shown in Figure 1.

**Figure 1:**
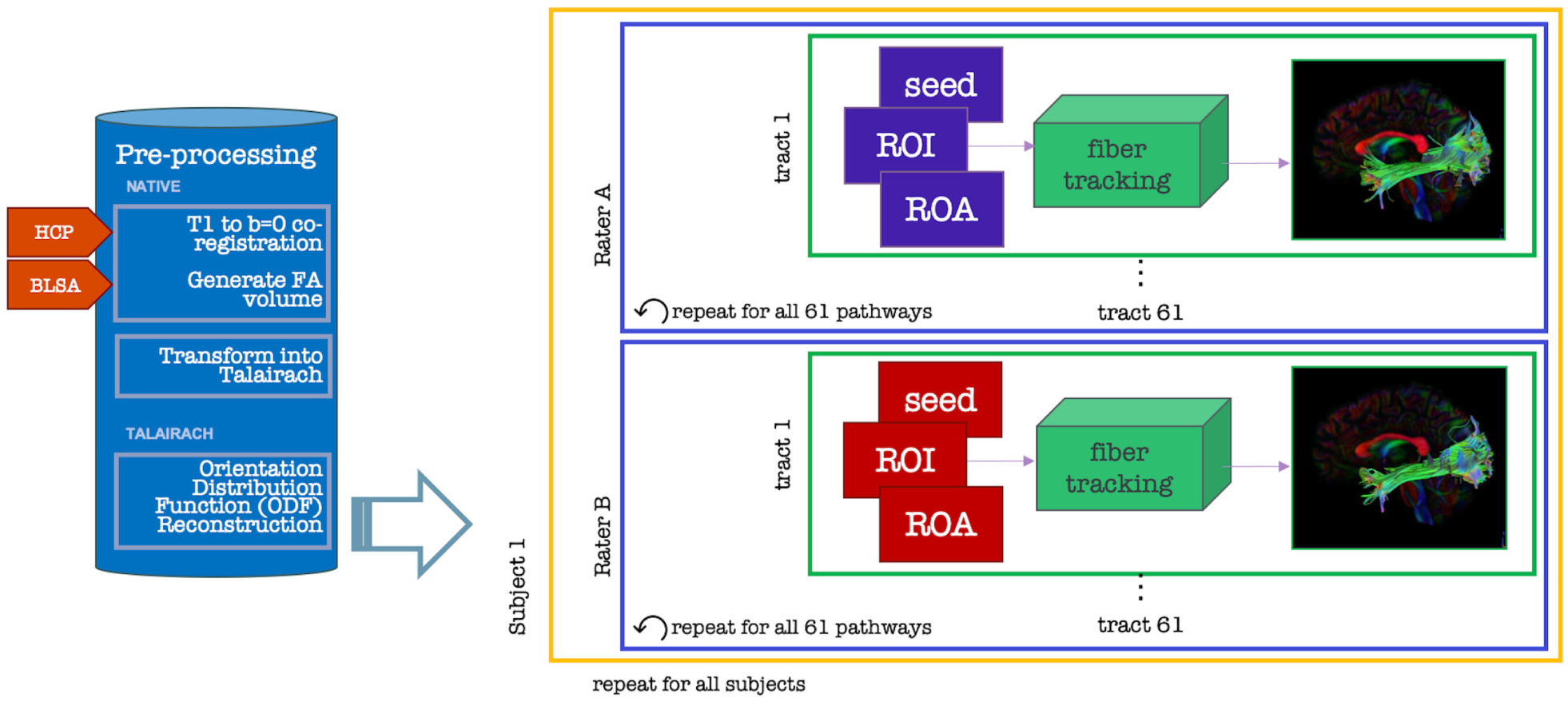
Pipeline overview: Pre-processing steps included susceptibility and eddy current corrections, and b0 signal normalization. A fractional anisotropy volume was created for every acquisition. After transferring each volume to Talairach space, 4th order spherical harmonic Q-ball imaging and generalized q-sampling are applied to BLSA and HCP datasets respectively to obtain orientation distribution functions, which are the inputs to DSI-Studio. The orientation distribution functions (ODFs) are used to create subject-specific 2D seed/ROI/ROA regions for each tract. Then region-based deterministic tractography is employed on the orientation maps. Each seed region was placed approximately in the middle of the bundle, all fibers entering the ROIs preserved and all passing through exclusion region (ROAs) masks are discarded. The results show two tractograms delineated by different raters.

The BLSA dataset used in this study contains 10 subjects with ages ranging between 57-77 years old (Ferrucci, 2008). The data were acquired after informed consent under institutional review board and accessed in de-identified form. Each session included a T1-weighted structural MP-RAGE (number of slices = 170, voxel size = 1.0×1.0×1.2mm3, reconstruction matrix = 256×256, flip angle = 8 degrees and TR/TE = 6.5ms/3.1ms) and two diffusion acquisitions. Diffusion data were acquired with a 3D spin-echo diffusion-weighted EPI sequence (TR = 7454 ms; TE = 75 ms). Each acquisition consisted of an initial b0 and 32 diffusion weighted volumes all with the same b-value of 700 s/mm2 (number of slices = 170, voxel size = 0.81×0.81×2.2mm3, reconstruction matrix=320×320, flip angle = 90 degrees, acquisition matrix=116×115, field of view = 260×260mm). Susceptibility correction (Andersson et al., 2003), eddy current (Andersson and Sotiropoulos, 2016) correction techniques, and b0 signal normalization were applied to the diffusion data as a preprocessing step.

The HCP dataset used in this study contains 10 healthy subjects with no known history of neuropathological and psychiatric diseases, ages ranging between 26-36. Diffusion scans obtained from HCP contained structural volume sampled at the same resolution as the diffusion data, the diffusion weighting (b-values) for each volume, the diffusion direction (b-vectors) for each volume, preprocessed diffusion time series, brain mask in diffusion space, and the effects of gradient nonlinearities on the b-values and bvectors for each voxel. Diffusion data were acquired with a 3D spin-echo diffusion weighted EPI sequence (TR = 5520 ms; TE = 89.50 ms). Each diffusion acquisition consisted of six b0 and 90 diffusion weighted volumes all with 3 shells of b=1000, 2000, and 3000s/mm2 interspersed with an approximately equal number of acquisitions on each shell within each run (number of slices = 111, voxel size = 1.25×1.25×1.25mm3, flip angle = 79 degrees, field of view = 210mm). Susceptibility correction (Andersson et al., 2003), eddy current (Andersson and Sotiropoulos, 2016) correction techniques, and b0 signal normalization were applied to the diffusion data as a preprocessing step.

### 2.2. DSI-Studio

DSI-Studio (http://dsi-studio.labsolver.org) is an end-user program for diffusion tensor image computation, region of interest ROI-based deterministic diffusion fiber tracking and 3D visualization. In this project, we have used DSI-Studio for both reconstruction and fiber tracking. Deterministic tractography is a maximum likelihood estimation approach for the fiber tracks, with high sensitivity parameters (high SNR, high anisotropy, small step size) generates reliable fiber pathway trajectories (Côté et al., 2013; Christidi et al., 2016). DSI-Studio adopts a generalized deterministic tracking algorithm that uses quantitative anisotropy (QA) to improve accuracy.

#### 2.2.1. Regions

DSI-Studio is an ROI-based tractography tool. Regions are defined as a set of voxels stored by their coordinates. There are different types of regions to inform fiber tracking. Multiple regions in combination can be used to control tracking when identifying fine grained details or complicated white matter structures. For example, if the tract is large, multiple regions of interest along with tract-specific exclusion regions can be used to delineate it. Regions can be manually drawn or created using a segmentation method. Example regions are shown in Figure 2. These regions are prior for the pathways, and extracted from the Eve atlas parcellation maps Figure 3.

**Figure 2:**
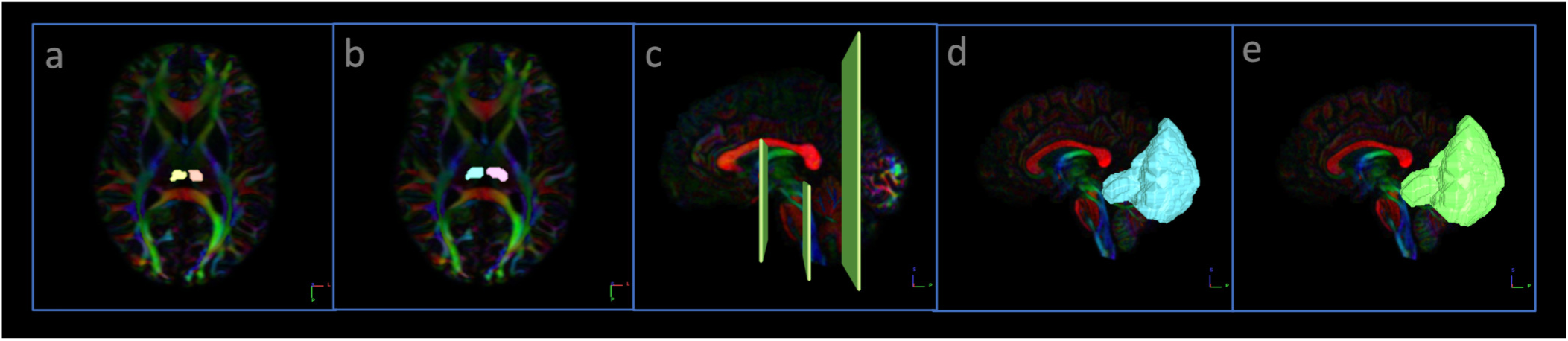
The regions for the region-based tractography that are used in TractEM are shown above (a) seed region, (b) ROI, (c), ROA, (d) 3D seed region, and (e) 3D end region (left to right).

**Figure 3:**
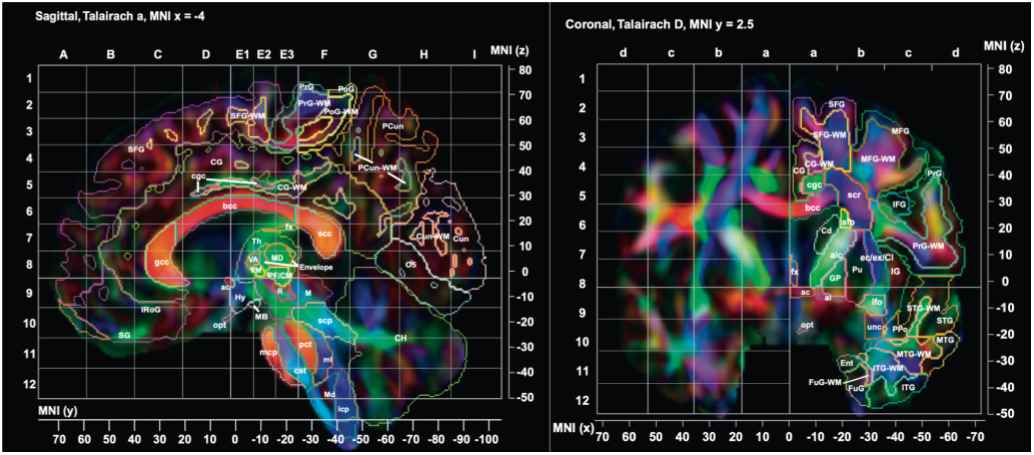
Region parcellations in Talairach from MRI atlas of human white matter used to develop TractEM protocol (a) coronal view of a random slice, (b) sagittal view of a near middle slice. Reprinted with the permission of Mori, S. et al., MRI atlas of human white matter, pg. 143 and pg. 249, Copyright Elsevier (2006).

##### Seed

Seed regions serve as starting points for the algorithm. A user labels seeding voxels which then are converted to starting points by the tracking algorithm. These starting points are at the subvoxel resolution to accommodate the algorithm. For example, a seed voxel placed at (1, 1, 1) will have subvoxel seeding placed within (0.5 1.5, 0.5 1.5, 0.5 1.5). Within the voxel region (0.5 1.5, 0.5 1.5, 0.5 1.5), DSI-Studio draws a point within the voxel range using a uniform distribution.

##### Region of interest (ROI)

These regions lead the fibers to where anatomical landmarks are. The streamlines are filtered in if passing through these landmarks, and eliminated from the tract if they are not.

##### Region of avoidance (ROA)

In the literature also commonly referred as NOT region or exclusion region. It is used to eliminate unwanted streamlines.

##### 3D Seed Region

For this project, subject-specific 3D seed regions were segmented by a multi-atlas method (Asman and Landman, 2013; Huo et al., 2016). 3D seed regions define the starting points for the four lobes.

##### 3D End Region

End region, as the name suggests, conserves only the tracts that end within a defined region. This is used for lobar region tracking. 3D end regions are created by dilating the multi-atlas segmented 3D seed regions for the same subject.

### 2.3. Pre-processing

The raw diffusion data were already corrected for head motion, eddy current effects, distortion (Van Essen et al., 2012), and b-table rotations (Schilling et al., 2019). When necessary T1 weighted images are co-registered to b=0. Using DSI-Studio, six elements tensor construction was performed on the diffusion-weighted image set to produce FA images in the subject’s native space. The FA map was registered to a Talairach template image (Talairach, 1988) using an affine transformation. The template image was generated by applying an affine transformation of Lancaster et al. (2007) to the probabilistic ICBM152-space FA template image from the FSL atlas distribution (Mori et al., 2005; Wakana et al., 2007; Hua et al., 2008). The affine transformation was applied to the diffusion-weighted images, and the appropriate corresponding adjustment was made to the b-vectors.

For orientation distribution function (ODF), two high angular resolution diffusion imaging (HARDI) models for voxel-wise reconstruction were selected, according to the image quality of each datasets. HARDI models are capable of characterizing multiple directions and resolving crossing fiber issues (Tuch, 1999; Tuch et al., 2002; Descoteaux et al., 2007).

In this implementation, for the BLSA dataset, we fit 4th order spherical harmonic Q-ball. QBI uses a single shell (single b-value) to estimate the dODF to represent the fiber orientation distribution. DSI-Studio adopts the spherical harmonics based QBI reconstruction method by Descoteaux et al. (2007). For multi shell HCP data, generalized Q-sampling (GQI) reconstruction was used. GQI in addition to being capable of resolving crossing fiber issues, can utilize multi-shell (multi-b-value) data and works best with fairly dense sampling of directions to similarly estimate the dODF (Yeh et al., 2010).

### 2.4. Protocol Design

The protocols have been designed with reference to the Eve protocol (Oishi et al., 2010) whenever possible. Eve is an effective guide for region drawing with high inter-rater reproducibility (k > 0.85 and CoV < 3 % (Mori et al., 2008)). A clear guide to a priori knowledge of the anatomy was prioritized when defining the TractEM protocols. The Eve atlas defines 50 white matter bundle parcellations slice by slice. Region-based tractography is performed using these regions of interest as prior. Example Eve white matter parcellation maps (WMPM) are shown in Figure 3. WMPM definitions are used to delineate tract-specific seed and ROI regions. A total of 61 unique pathways were selected such that most of the brain was accounted for: 53 of these pathways, 22 of which have bilateral pairs of homologous tracts (22×2=44), 5 commissural pathways, 3 of the brainstem tracts and the optic radiation were manually identified with Eve. In addition to the Eve-defined pathways, 8 subject-specific lobar regions were defined by a multi-atlas segmentation algorithm (Huo et al., 2016).

During the design and the development of the protocols, the high-acquisition (multi-shell) HCP dataset was used. Contrary to the Eve atlas, TractEM protocols do not divide anatomical fibers into multiple regions. Instead, anatomically consistent regions are merged into individual tracts. For the 53 Eve-defined pathways, seed regions were drawn roughly at the center of each tract. For bilateral tracts, seed regions were placed in both hemispheres. As white matter structural complexity increased, we increased the number of additional ROIs and ROAs in order to capture a tract as consistent as possible with its anatomical definition. In this manuscript, the complexity of the protocols is referred to as simple and complex tracts. If a protocol contains only a single seed region, e.g. genu of the corpus callosum (GCC), we refer it as a simple tract. If a protocol includes multiple seed/ROI regions in combination, this is referred as a complex tract, e.g. inferior longitudinal fasciculus (ILF) (Please see Figure 4). We further divided the complex tract labels as 2-unique and 3+ unique labels. For the lobar regions, multi-atlas segmented subject-specific 3D lobar masks were used as seed regions and 3x-dilated lobar masks were used as end regions. ROA regions were guided in the protocols in order to isolate the tract or region of interest to fit into its anatomical definition for both the pathways and the lobes.

**Figure 4:**
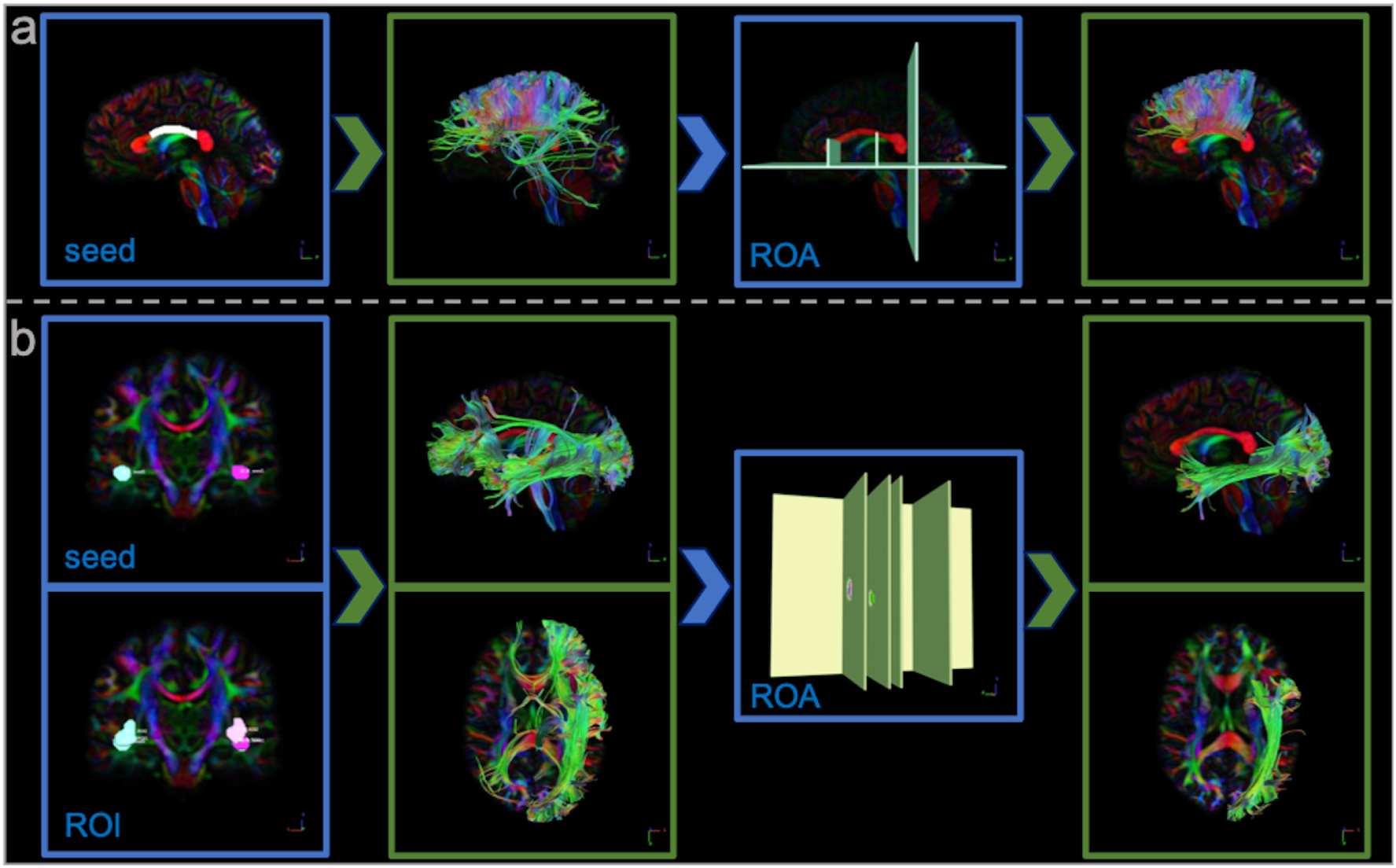
The application of TractEM on a simple (a) and a complex (b) tract. (a) Body of the corpus callosum is traced with a single seed region on the sagittal axis and three ROA regions one on the axial, three on the coronal axes, (b) bilateral inferior longitudinal fasciculus is traced using one seed, two ROI regions for each side and multiple ROA regions.

### 2.5. Application

#### 2.5.1. Tracking Parameters

Tracking parameters were chosen carefully to reduce the number of false positive fibers and to ensure that the tracking does not suffer from premature termination. Parameters were termination index normalized QA (nqa) with threshold of 0.1 is used to ensure the maximum number of fibers are produced, smoothing of 1, tract count of 100K, track length constraints 15-300 mm for lobar regions, and 30-300mm for the rest of the tracts. The QA threshold is the suggested minimum as lower values may produce false fibers (http://dsi-studio.labsolver.org). Angular and QA thresholds are the determining parameters of how the fibers are tracked from the region of interest. Angular threshold is set to zero, therefore stopping criteria is not determined by this parameter; instead the length constraint or end regions serve as an end condition. The tracking algorithm is streamline (Euler), seed orientation is primary, and the direction interpolation is set to trilinear. Randomize seeding is off. This parameter sets the random seed generator to a constant to ensure tracking results are identical, and reproducible.

#### 2.5.2. Fiber Tracking with DSI-Studio

The TractEM project (https://my.vanderbilt.edu/tractem) contains 35 tract specific protocols. Figure 4(a) visualizes application of the protocol for a simple (top) and complex (bottom) track. The corpus callosum is an example of a simple track. Here, a sagittal seed region (at approx. sagittal slice 78) was created. The anterior extent of the seed region lines up with the most posterior part of the genu, whereas the posterior extent lines up with the most anterior part of the splenium. Then fiber tracking was executed. Based on the tracking output, 2-4 ROA regions were placed with one on an axial slice inferior to the seed regions, one on a coronal slice posterior to the seed region and when necessary a couple more on coronal slices inferior to the seed regions to control leakage into the fornix area. A second fiber tracking was then performed. The output was stored as streamlines (.trk) and converted into tract density maps (.nii). Figure 4(b) depicts the protocol for inferior longitudinal fasciculus, considered a complex pathway, which requires multiple seed, ROI, and ROA regions to capture the pathway. Lastly, lobar regions were traced using 3D subject-specific lobar masks created by a multi-atlas segmentation algorithm.

#### 2.5.3. Data Partitioning, Submission Issues and Repeats

The TractEM protocols are developed on 10 subjects from the high acquisition HCP dataset. The reproducibility testing of the protocols was made with an additional 10 subjects from the BLSA dataset, totaling 20 subjects. In order to test the reproducibility of the protocols, every subject has been traced at least two times by different raters.

The raters submitted their tract-specific labels along with the streamlines and the density images of these pathways. Raters were allowed to use any version of DSI-Studio. After all the data was gathered, to reduce the variability due to DSI-Studio version differences and to ensure that the data was free of human errors (naming etc.), we reran the protocols for all subjects using the manual labels. Auto-tracking was capped to 15 minutes. If the auto-tracking did not result in the same streamline submitted, the tracts were manually re-labeled by another rater and the missing data has been replaced. Some errors due to human fallibility was left unfixed in the dataset. Comprehensive list of these issues as follows: (1) some raters sought placement guidance from other raters, resulting in very similar labels, (2) some raters used the same labels for different subjects, (3) some raters did not closely follow the protocols, (4) some raters traced the same-subject tract more than once.

## 3. Results

With the TractEM, we designed fast-tracking protocols that are driven by state-of-the-art tractography. Using the TractEM, we created a dataset that contains 2562 manually identified tracts. Fiber tracking for one subject took less than 6 hours with no neuroanatomical expertise required. This time included learning how to do fiber tracking following the protocols. Evaluation of the protocol was performed in 20 subjects where each subject was traced at least twice by different raters and reproducibility was assessed both qualitatively and quantitatively.

This section describes the evaluation procedure throughout the development and validation of the protocols. It is organized as follows. First, quality assessments were administered to check the validity of the tracts and approve/reject the methodology for the construction of tracts. Second, qualitative analyses demonstrated the resulting fiber tracts. Concluding analyses consisted of measures of inter-rater tract reliability, inter-subject interrater variability, and intra-subject rater reproducibility.

### 3.1. Quality Assessment

Quality assessment is a crucial step to ensure validity of tracts. We have prepared a performance analysis for immediate rater feedback. An automated pdf report is generated for each tract. It includes tract density visualization overlaid on the subject’s FA images for all three views: sagittal, coronal, axial, and a 3D representation of the streamline Figure 5. This tool served as a quick visual sanity check to see if the assembled tract appears is consistent with the expected morphology. Initially, we used this assessment to improve the protocol. Its secondary and more important use is to check if the rater interpreted the protocol correctly. This tool served as a quick visual sanity check to see if the assembled tract appears consistent with the expected morphology. Initially, we used this assessment to improve the protocol. Its secondary use is to check if the rater interpreted the protocol correctly.

**Figure 5:**
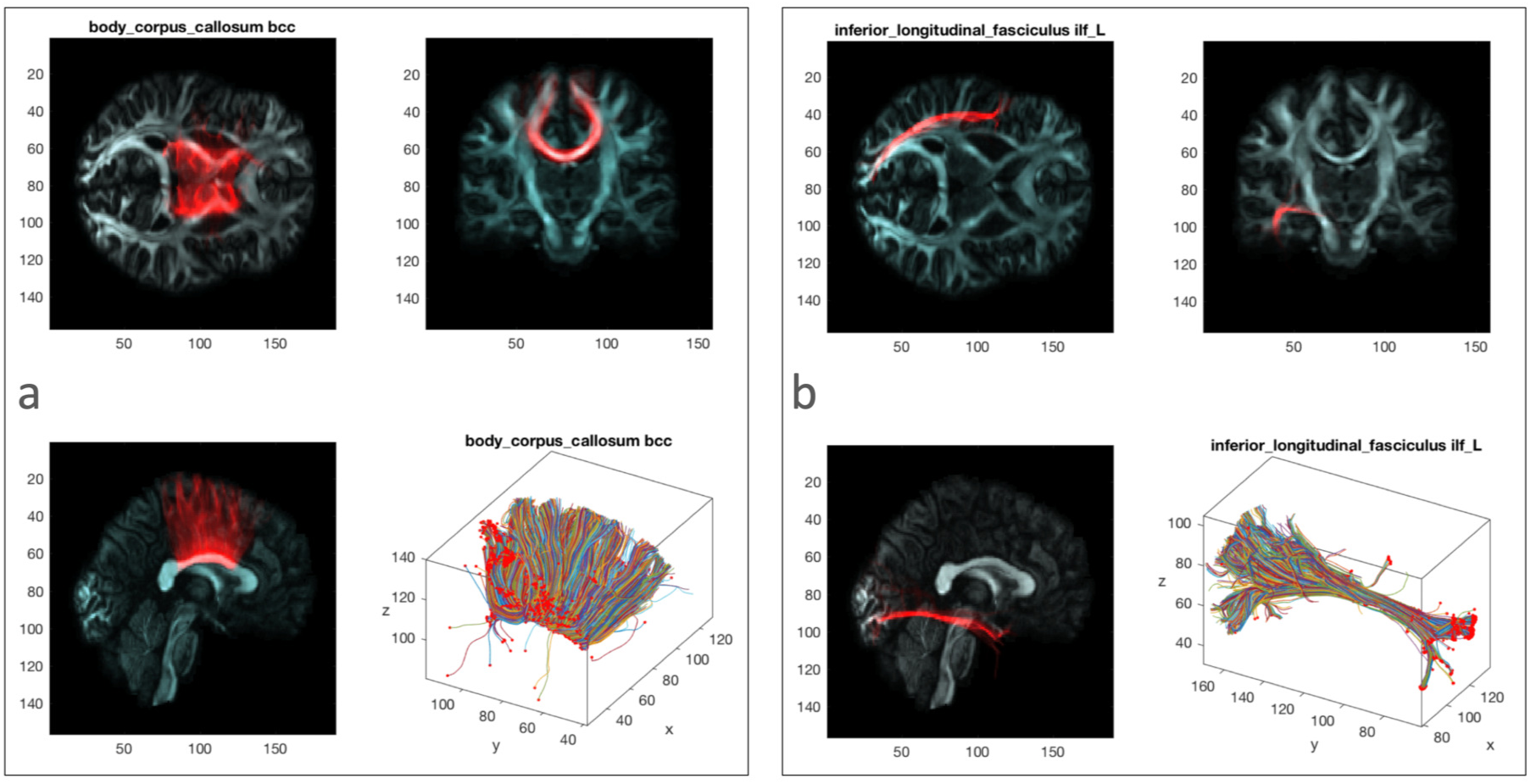
Quality Assessment PDF: Axial, coronal, sagittal view of the density maps, and 3D visualization of streamlines for (a) body of the corpus callosum, and (b) inferior longitudinal fasciculus.

### 3.2. Qualitative Analysis

Qualitative analysis is vital in order to contextualize the estimated fiber bundles. The characterization of individual fiber bundles can be inferred from these images, and the tracts can be investigated by their behavior (i.e. if the tract splits off from, is close to, integrates into another tract). Figure B.1 shows the resulting averaged density maps of 61 white matter tracts for 20 subjects reconstructed using TractEM. Inter-regional relations and connections across the brain can be seen from these 3D fine detailed parcellation maps. Information about spatial location, shape, tract volume, proximity to the other tracts are important bases when assessing reproducibility. Thus, we also provide the tract visualizations in five structural categories (1) tracts in the brainstem (Figure 6), (2) projection fibers which connect cerebral cortex to the central regions and to the other end to spinal cord (Figure 7), (3) association fibers which connect different region of cerebral cortex (Figure 8), (4) commissural fibers which connect the two hemispheres (Figure 9), and (5) lobes which are the four main cortical divisions (Figure 10).

**Figure 6:**
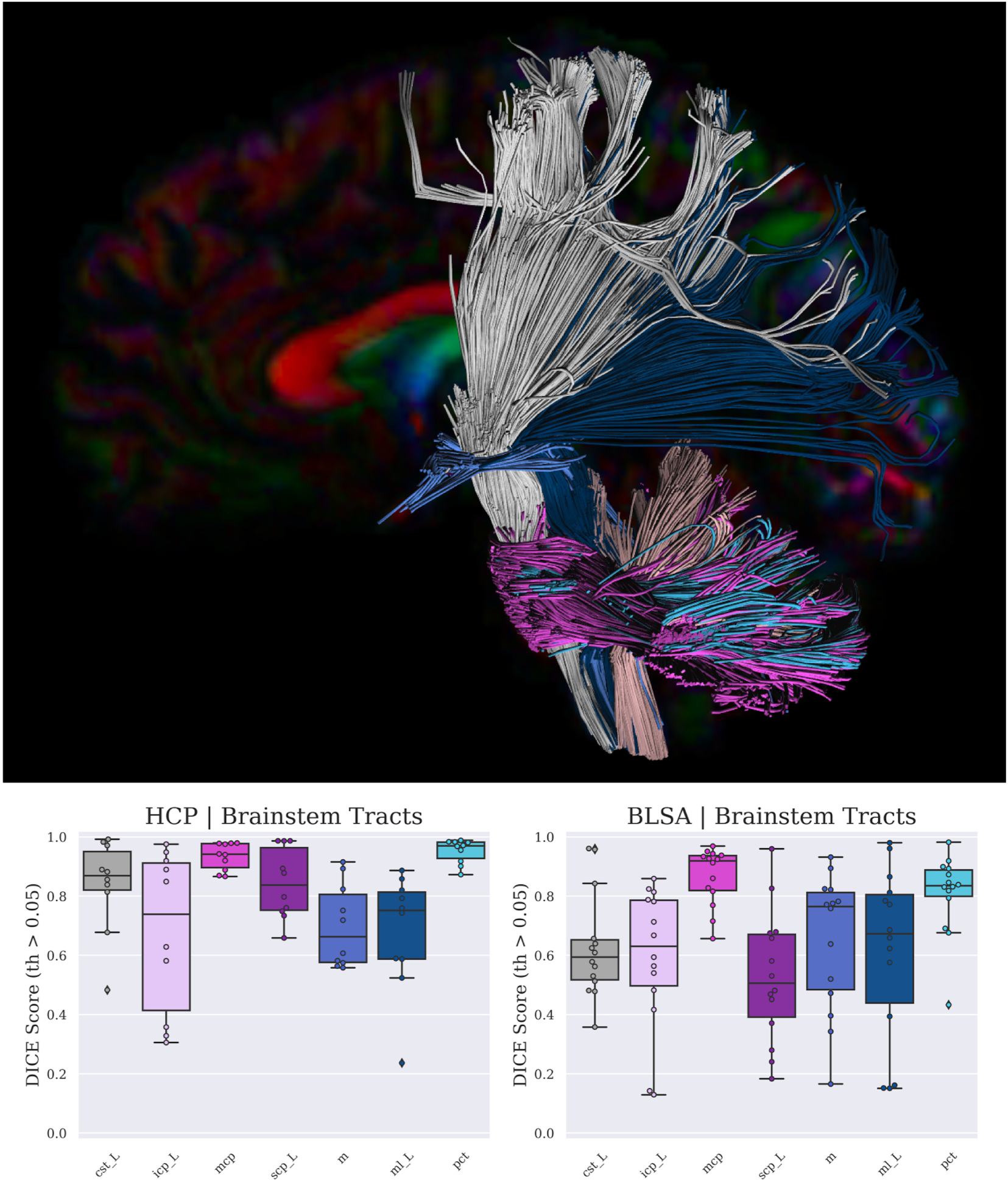
Visual representation of the TractEM definitons (above), inter-rater **brainstem** tract reliability both for HCP and BLSA datasets (below); corticospinal tract left (cst_L), inferior cerebellar peduncle left (icp_L), middle cerebellar peduncle (mcp), superior cerebellar peduncle left (scp_L), midbrain (m), medial lemniscus left (ml_L) and pontine crossing tract left (pct_L).

**Figure 7:**
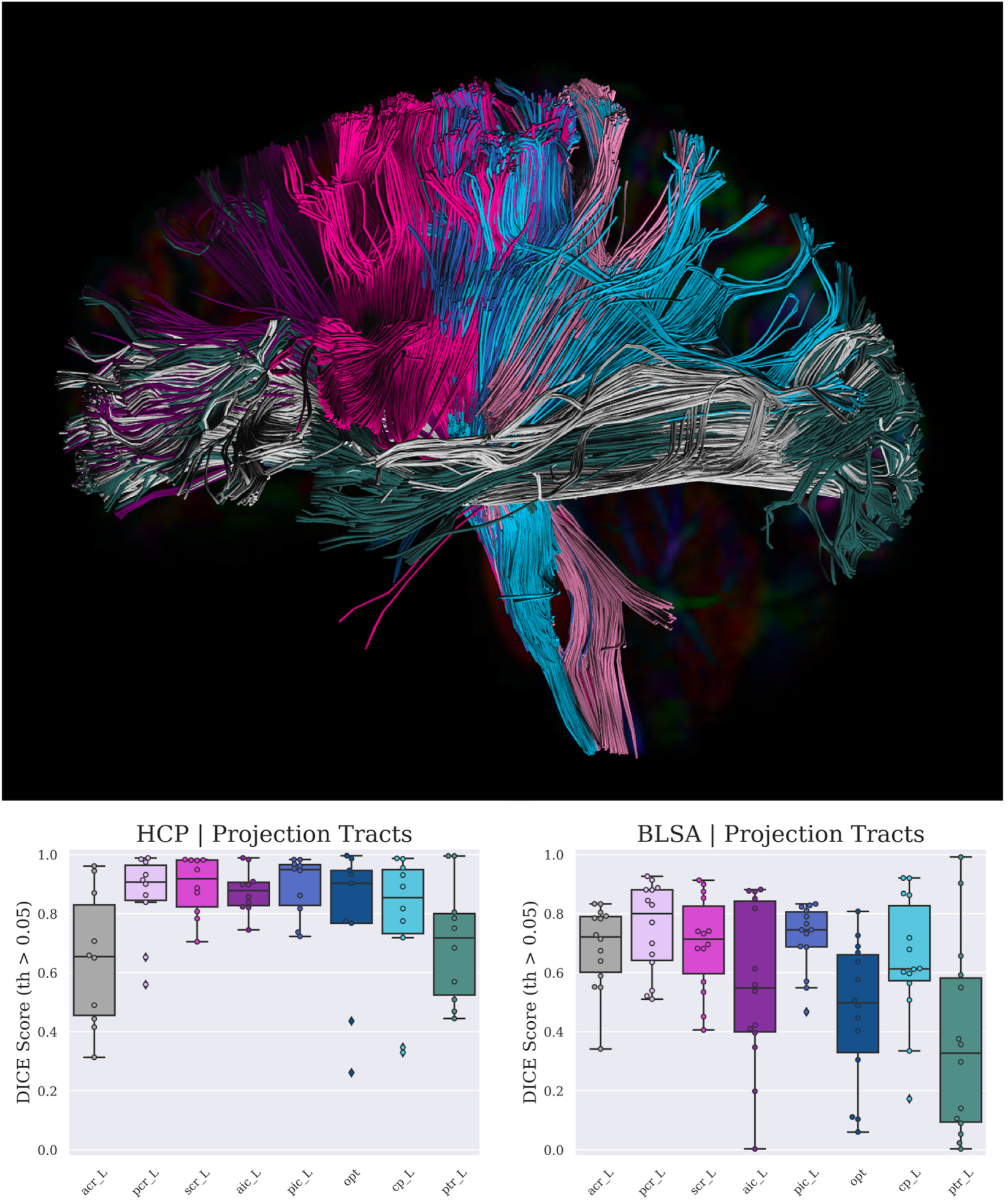
Visual representation of the TractEM definitions (above), inter-rater **projection** tract reliability both for HCP and BLSA datasets (below); anterior corona radiata left (acr_L), posterior corona radiata left (pcr_L), superior corona radiata left (scr_L), anterior limb of internal capsule left (aic_L), posterior limb of internal capsule left (pic_L), optic radiation (opt), cerebral peduncle left (cp_L) and posterior thalamic radiation left (ptr_L).

**Figure 8:**
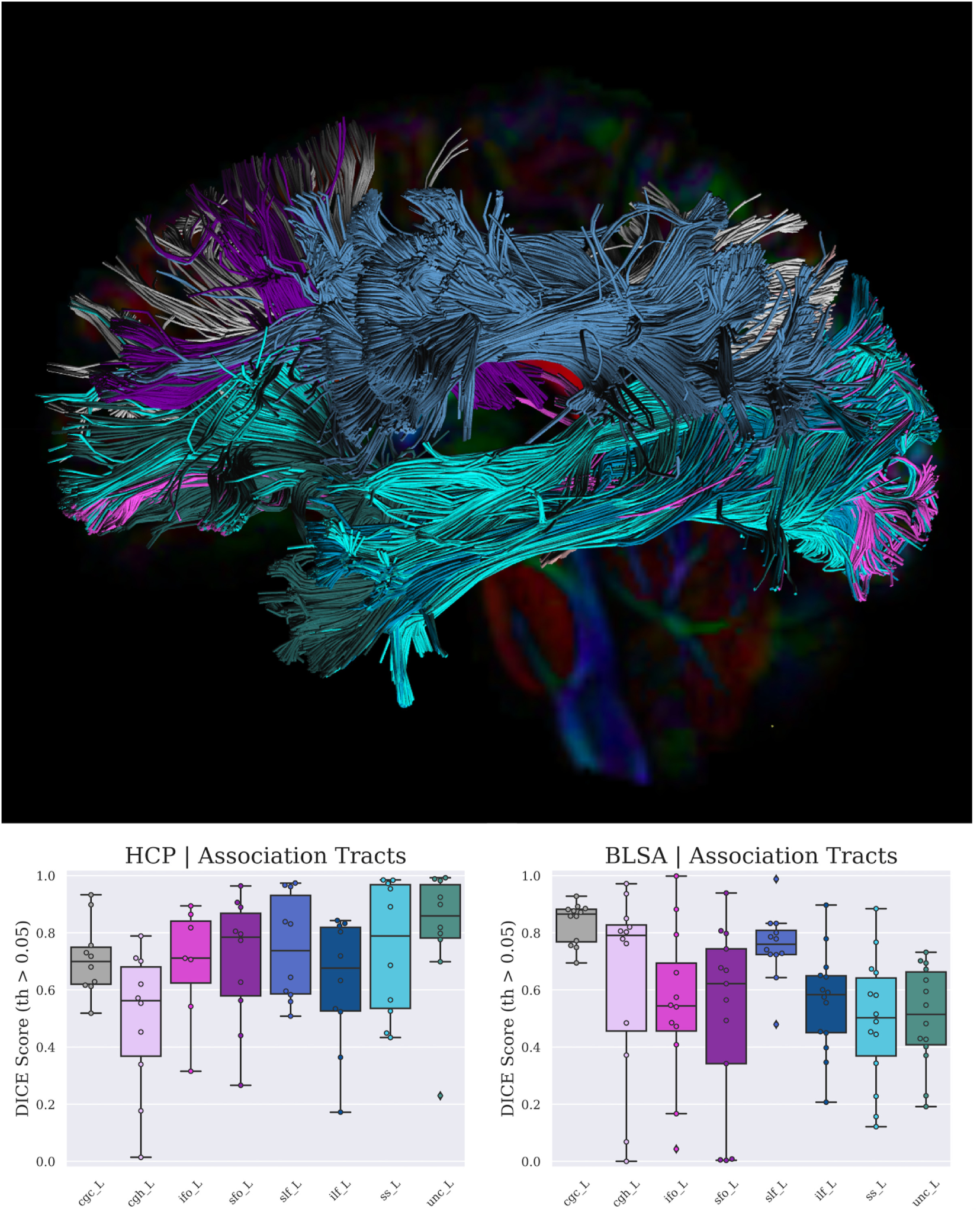
Visual representation of the TractEM definitions (above), inter-rater **association** tract reliability both for HCP and BLSA datasets (below); cingulum cingulate gyrus left (cgc_L), cingulum hippocampal left (cgh_L), inferior fronto-occipital fasciculus left (ifo_L), superior fronto-occipital fasciculus left (sfo_L), superior longitudinal fasciculus left (slf_L), inferior longitudinal fasciculus left (ilf_L), sagittal stratum left (ss_L) and uncinate fasciculus left (unc_L).

**Figure 9:**
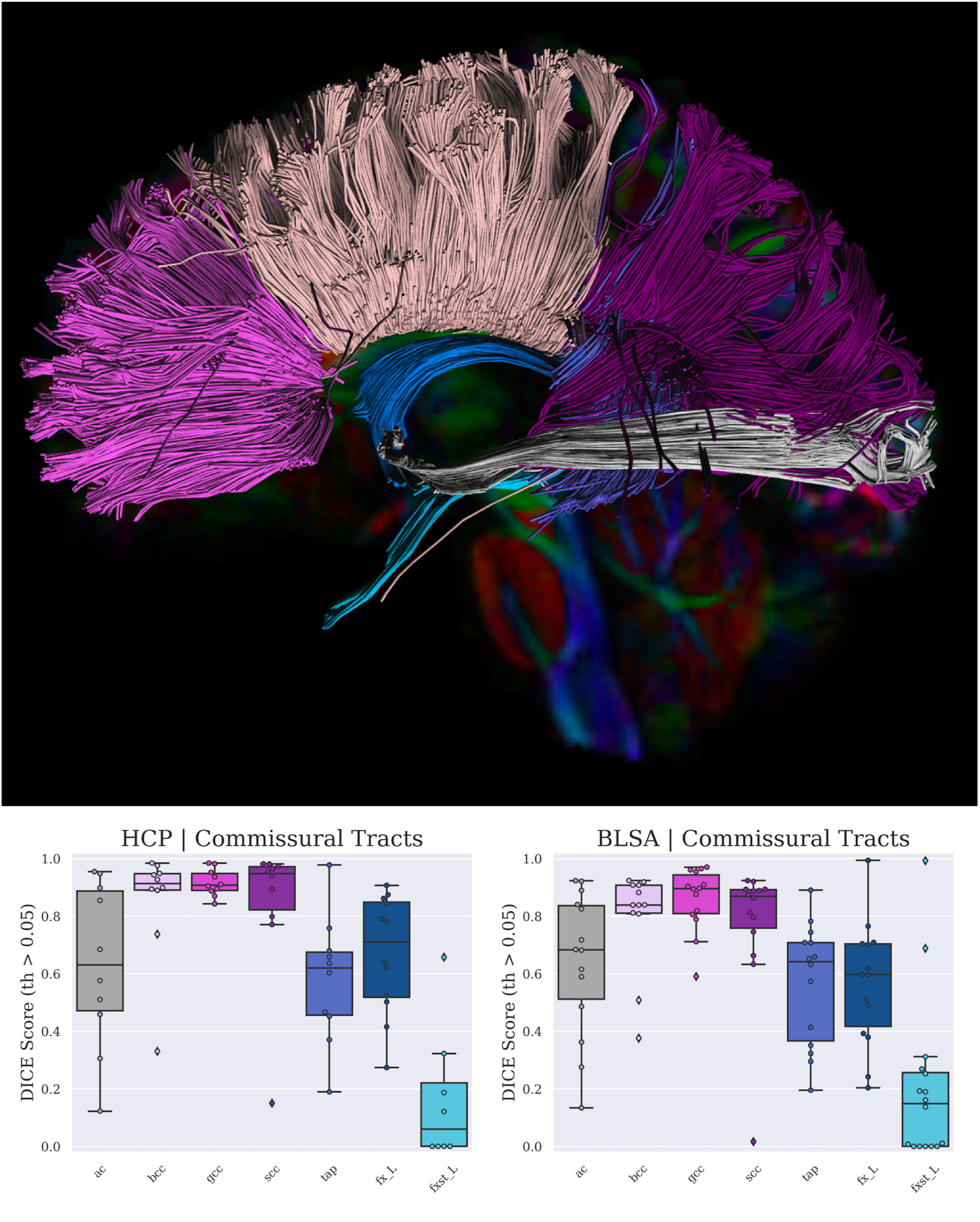
Visual representation of the TractEM definitions (above), inter-rater **commissural** tract reliability both for HCP and BLSA datasets (below); anterior commissure (ac), body of the corpus callosum (bcc), genu of the corpus callosum (gcc), splenium of the corpus callosum (scc), tapetum (tap), fornix left (fx_L) and fornix stria terminalis left (fxst_L).

**Figure 10:**
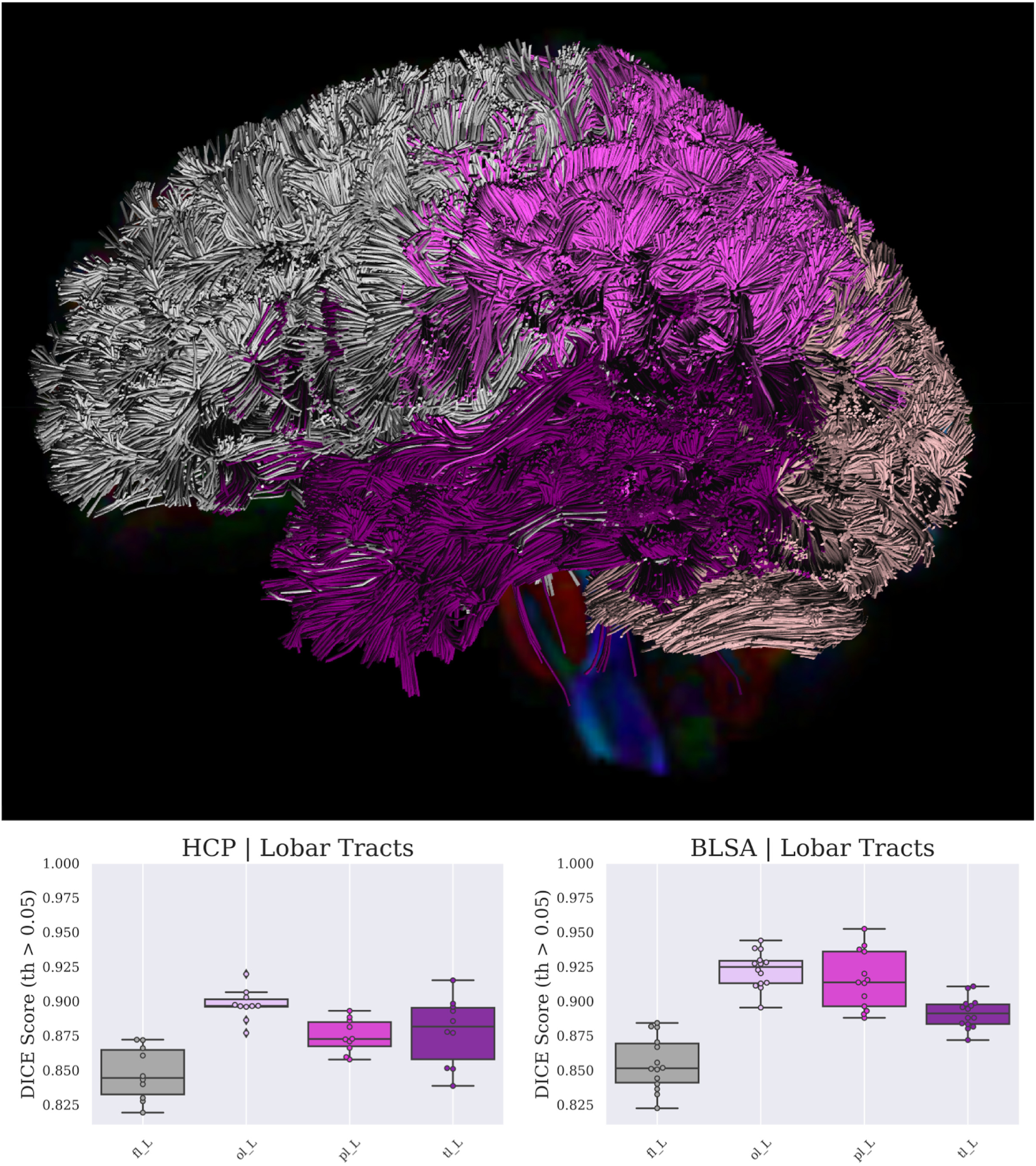
Visual representation of the TractEM definitions (above), inter-rater **lobar** tract reliability both for HCP and BLSA datasets (below); frontal lobe (fl_L), parietal lobe (pl_L), occipital lobe (ol_L) and temporal lobe (tl_L).

### 3.3. Quantitative Analysis

Reproducibility analysis of the protocols is made possible by having white matter tracts of each subject traced by two separate raters. The protocols have been tested by ten raters who have no prior experience in tractography. Reproducibility was assessed between raters and across tracts. To measure the reliability of the protocols, we analyzed inter-rater tract reliability. We also assessed inter-subject variability to highlight anatomical differences between subjects and intra-subject rater reproducibility to confirm the robustness of the protocol.

#### 3.3.1. Inter-rater Tract Reliability

Same subject analysis leverages reproducibility analysis of a protocol by removing the anatomical variation. Intra-subject inter-rater tract reproducibility assessment was made by evaluating rater agreement for each tract. The score was calculated by averaging similarity across all subjects for a corresponding tract. DICE correlation coefficient (DICE), intra-class correlation (ICC) and root mean squared error (rMSE) metrics (see metrics in details in Appendix A) were used for the analysis. For each tract, medians and standard deviations were plotted for both datasets. HCP has shown DICE (threshold > 0.05) > 0.7, ICC > 0.8 reproducibility on most tracts. BLSA results have shown DICE (threshold > 0.05) > 0.6, ICC > 0.6 reproducibility for most tracts, which is lower than the HCP dataset. We defined the pathways during the TractEM development stage as lobar regions, simple and complex tracts. This description simply explains the type and number of regions used to track these pathways. Lobar regions were created using a multi-atlas segmentation method, simple tract protocols have one seed, and complex tract protocols are defined with the combination of 1+ seed and/or ROI regions. Lobar regions exhibit very high reproducibility across both datasets. Simple tracts showed generally higher reproducibility than more complex tracts. For simple tracts, even though HCP showed a more distinguishable division of reproducibility between simple and complex tracts, the BLSA dataset shows the same trend. The reproducibility for simple tracts in the HCP, DICE scores (th > 0.05) ranged from very reproducible 0.95 to highly reproducible 0.68 and ICC scores fall into a similar scale between 0.98-0.75; for the BLSA, 0.92 > DICE (th > 0.05) > 0.53, 0.96 > ICC > 0.49. The reproducibility of tracts with two or more unique labels in the HCP, DICE (th > 0.05) scores ranged between 0.97 and 0.4 and ICC scores ranged between 0.99 and 0.47; for the BLSA, 0.88 > DICE (th = 0.05) > 0.1 and 0.93 > ICC > 0.13.

The distribution of these tracts is detailed for the 61 unique tracts and for each error metric, it is shown in the form of box plots Appendix B, Figure B.2-9.

Inter-hemisphere asymmetry was observed even though the protocols were the same for the same tract. Bilaterality of white matter pathways has been reported in many previous studies in healthy subjects (Galaburda et al., 1978; Pujol et al., 2002; Park et al., 2004; Kanaan et al., 2012). The asymmetry suggested to occur during development, with age, in disease as well as it is observed between genders (Toga and Thompson, 2003). Bilateral reproducibility similarly varied more when the tract fell into the complex tract category.

#### 3.3.2. Inter-subject Variability

Inter-subject variability was analyzed to understand anatomical variability across subjects. The variability is calculated by measuring the similarity between every subject for each dataset. To note that these analyses also have the variability introduced from inter-rater reproducibility. Inter-subject variability is a crucial contributor to the reproducibility analyses of the tracts. This information is provided to aid a fair evaluation of the intra-subject tract reliability. The results have confirmed the variability (0.6 > DICE, ICC) that rises from microstructural differences between subjects shown in Appendix B, Figure B.10-13.

#### 3.3.3. Intra-subject Rater Reproducibility

The rater reproducibility evaluates the performance of different raters on images acquired under different acquisition parameters. A single rater tested all 35 protocols on their assigned subjects. Depending on their availability, the raters traced 1-5 subjects per person on each dataset (Appendix B, Table B.1). On unanticipated occurrences, some raters traced the same-subject tract more than once. This issue was raised from failures in following the protocols. 67 tracts on BLSA data and only 2 tracts on HCP data were replaced and resulted in intra-rater pairs. These tracts were also included in the tract reliability analyses. Before the study, raters had neuroanatomical backgrounds ranging from no-knowledge to basic-knowledge. The results have shown homogeneity thus confirming the robustness of the protocols. For the multi-shell HCP dataset, the rater reproducibility was expectedly higher than the single-shell BLSA dataset. The overall rater reproducibility varied by less than 25% within each dataset. Evaluations across the five metrics are shown for both BLSA and HCP datasets in Appendix B, Figure B.14-15.

## 4. General Discussion

The major advantage of diffusion tractography is that it provides reliable correspondence with white matter fiber pathways in vivo. Qualitative and quantitative identification of the estimated fiber pathways are crucial for the veracity of connectivity studies that rely on manual or automated approaches. Reproducible tractography can provide a backbone to the group-wise analyses and longitudinal testing studies for which a comprehensive modern atlas is essential. Despite the importance, there are a limited number of reproducibility studies with limited number of fiber pathways (Heiervang et al., 2006; Wakana et al., 2007; Besseling et al., 2012).

With the TractEM project, we tested 61 unique tracts using 35 protocols. The most reproducible fiber bundles were the lobar tracts, followed by corpus callosum (CC), cingulum, cerebral peduncle (CP), middle cerebral peduncle (MCP), corticospinal tract (CST), superior longitudinal fasciculus (SLF), inferior longitudinal fasciculus (ILF), inferior fronto-occipital fasciculus (IFO), and uncinate fasciculus (UNC) on both datasets. In contrast, FX, fornix stria-terminalis (FXST), superior cerebellar peduncle (SCP), posterior thalamic radiation (PTR) exhibited low reproducibility. Tracking sometimes resulted in less than 100K streamlines (tract count). FXST takes a particularly long time, results in lower tract count. Anecdotally, reproducibility could be increased if more time allowed. The low reproducibility tracts generally fitted in our complex tract description (please refer to Section 2.4). It can be inferred that as the complexity of manual protocols increases, the rater reproducibility decreases. In addition to the complexity, manual region drawing and slice selection affected reproducibility. This reproducibility issue progressively improved as the protocol evolved by selecting easily distinguishable parcels (from the Eve-parcellation maps) and adding specificity to the protocol descriptions (i.e. approximated slice numbers).

As mentioned earlier, we assessed reproducibility on probabilistic density images. A direct comparison of the reproducibility could not be made between our analyses and ones in the literature due to the selection of different modalities and metrics. One of the earliest works on reproducible tractography is from Wakana et al. (2007) where they performed reproducibility analyses using kappa measure on FA and T2 for 11 tracts, 8 of which exist in TractEM (CGC, CGH, CST, SLF, ILF, IFO and UNC). The inter-institution inter-rater reproducibility is the closest assessment to our evaluation of tract reproducibility. These assessments resulted in average correlation ranging between 0.73-0.95 for the 8 tracts evaluated (Wakana et al., 2007). Heiervang et al. (2006) tested their most reproducible method of seed definition for probabilistic tractography. They evaluated tract volume overlap for only three TractEM corresponding tracts: cingulum bundle, GCC and optic radiation. Their results have shown inter-session and inter-subject mean overlap 81% and 50% respectively. Catani and De Schotten (2008) suggested a template guide for some of the association, projection and commissural pathways. They investigated cingulum, ILF, UNC, IFO, CC, AC, fornix (FX) and cerebral tracts and presented qualitative representations of them. The anatomical references are in agreement with our protocols. Another group, Besseling et al. (2012), did tract morphology similarity analyses for cingulum and GCC. They assessed reproducibility for proximal (excluding cortical projections) and extended (including cortical projections) streamlines.. The reproducibility of these two tracts was measured using DICE with a threshold of 5%. For the extended representation of cingulum and GCC, average DICE values were 0.82 and 0.84 respectively.

Different automated protocols are capable of producing streamlines, highly reproducible even for small pathways, and accounts for the whole brain (Garyfallidis et al., 2018; Wasserthal et al., 2018; Yeh et al., 2018).Nonetheless, these methods rely on manual delineations as their prior. This need alone highlights the importance of reproducibility studies at the manual level. Tract specific protocols in combination with subject-specific delineations can provide a backbone to automated methods.

We have traced and analyzed lateral and single tracts that co-exist in both hemispheres. Our results quantitatively confirm the inter-hemisphere asymmetry that has been shown in multiple studies (Toga and Thompson, 2003; Wakana et al., 2007; de Schotten et al., 2011) (see Appendix B, Figure S.2-9). It was not clear what caused this asymmetry, whether it was anisotropy or the size of the tracts. From a broader perspective, lateralization is evolutionary, hereditary, and developmental, and may occur as the result of disease. Mapping the asymmetry between hemispheres may be a very useful tool to identify abnormalities and the relationship between neighboring white matter tracts.

TractEM allows non-expert users to create whole brain tractography datasets. At its initial developmental stage, it has shown encouraging results. Using the protocols constructing a full dataset with 35 tracts with their bilateral counterparts is feasible (< 6 h/subject).

## 5. Conclusions

In this study, we addressed the lack of available manual labeling protocols that are reproducible in spite of human subjectivity. In response to this challenge, we present the TractEM protocols. The TractEM project allows expert/non-expert users to create and curate reproducible pathways across the brain and can be applied generally to large cohorts. To the best of our knowledge, this is the first study that assesses human variability across minor and major pathways on both healthy and aging study data.

## Data and Code Availability

TractEM is an open source project, in terms of data and labeling guidelines. Comments and discussion are open for each tract. The frameworks enable versioning of tract definitions as consensus definitions evolve. By crowd sourcing our results, we aim to obtain expert validation for each tract. The raw input data, Talairach-aligned data, DSI-Studio-ready input data, and TractEM results are made available on our website, https://my.vanderbilt.edu/tractem.

## Acknowledgements

This work was supported by the National Institutes of Health under award numbers R01EB017230, T32EB001628, and in part by ViSE/VICTR VR3029 and the National Center for Research Resources, Grant UL1 RR024975-01. This research was conducted with the support from Intramural Research Program, National Institute on Aging, NIH. The content is solely the responsibility of the authors and does not necessarily represent the official views of the NIH.

## Appendix A. Metrics For Quantitative Analysis

We assessed reproducibility to test the TractEM protocols using DICE correlation coefficient, intra-class correlation and root mean squared error. Reproducibility analysis is essential to validation of robustness. The reader should note that since we did not acquire ground truth for this study for every intra-subject comparison, one rater has randomly chosen from the pair and treated as ground truth.

### Dice Correlation Coefficient

Dice correlation coefficient is a volumetric overlap metric. It quantifies binarized morphological similarity. Dice coefficient is defined as

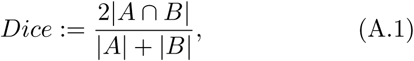

where A and B represent binary images with values “0” and “1” for each voxel and computed as follows:

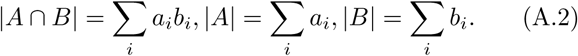

### Intraclass Correlation

The interclass correlation coefficient (ICC) (McGraw and Wong, 1996) is a measure of correlations between pairs of observations that do not necessarily have an order, or have obvious labels. It is common to use the ICC as a measure of agreement among observers; in our case it is used as a measure of agreement between two segmentations. Inter-class correlation is defined as

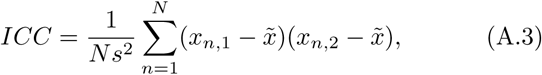

where

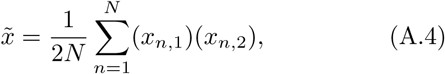

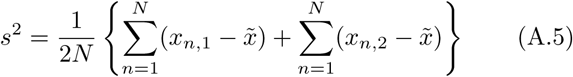

## Appendix B. Qualitative and Quantitative Representations of Results

This section is reserved for supplemental figures.

**Table B.1:** Number of tracts labeled by each rater are shown in the table for each dataset. The number in each box indicates the number of total usable tracts that the raters traced. Each subject was traced at least 2 times for reproducibility analysis. For 10 subjects 35 protocols, this number should add up to total 700 (=10×2×35) minimum number of pathways per dataset. The last column shows the after math. For the BLSA dataset, total of 8 raters and for the HCP dataset, total of 9 raters tested the protocols.

| **Rater ID**<br>**Dataset** | R1 | R2 | R3 | R4 | R5 | R6 | R7 | R8 | R9 | R10 | Total |
| --- | --- | --- | --- | --- | --- | --- | --- | --- | --- | --- | --- |
| BLSA | 87 | 95 | - | 177 | 101 | 64 | 8 | 75 | 57 | 104 | 768 |
| HCP | 55 | 68 | 35 | 104 | 104 | 20 | 39 | 78 | 99 | 104 | 706 |

**Figure B.1:**
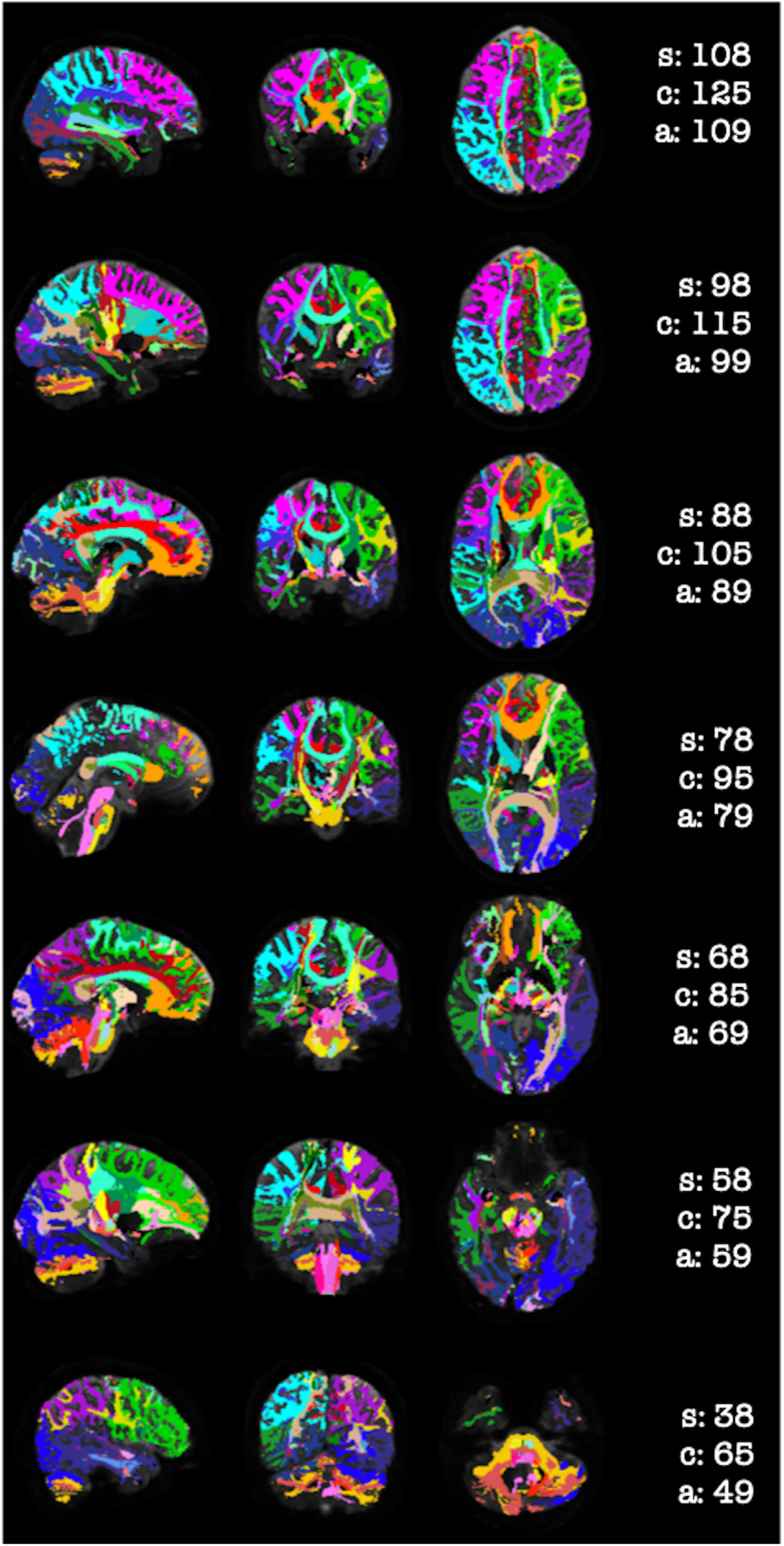
Tractography-based white matter atlas including 10 subjects total of 61 tracts with bilateral pairs is shown above from different slice views for all three views, sagittal (s), coronal (c), and axial (a), with slice numbers indicated.

**Figure B.2:**
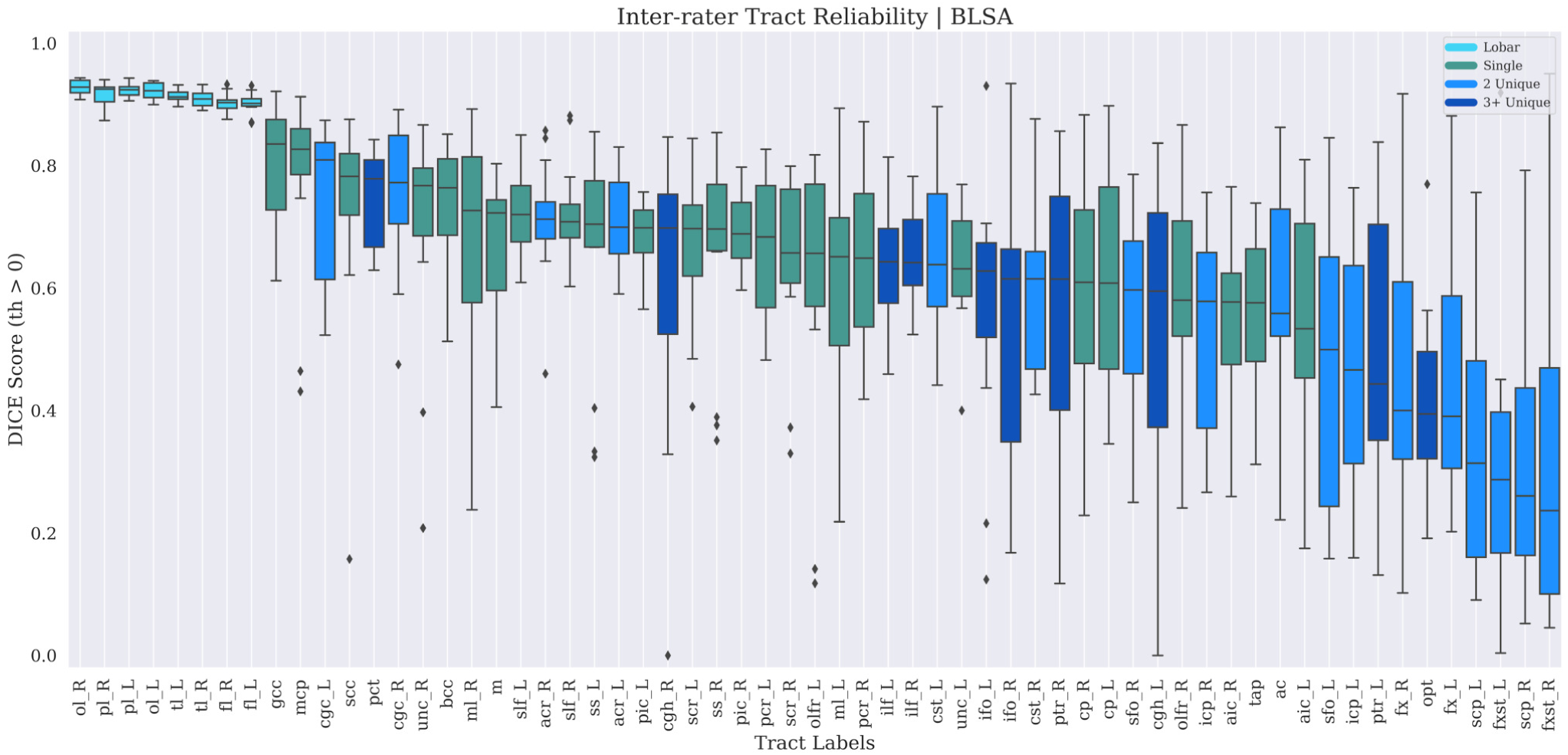
TractEM tract reliability for all 61 fiber bundles is shown. Density images were binarized with a threshold > 0 for similarity analysis and similarity was calculated using **DICE correlation coefficient (DICE)**. Each boxplot represents the similarity of same-subject different-rater pairs for 10 subjects from the BLSA dataset (see Section 3.3 Inter-rater Tract Reliability for the exceptions). The tracts are sorted from high to low reproducibility. Lobar regions are demonstrated with turquoise color, single label protocols are shown in dark green, protocols composed of 2-unique labels are represented by blue and protocols composed of 3+ unique labels are marked with dark blue color (see Section 2.4 TractEM Protocol Design for tract group descriptions).

**Figure B.3:**
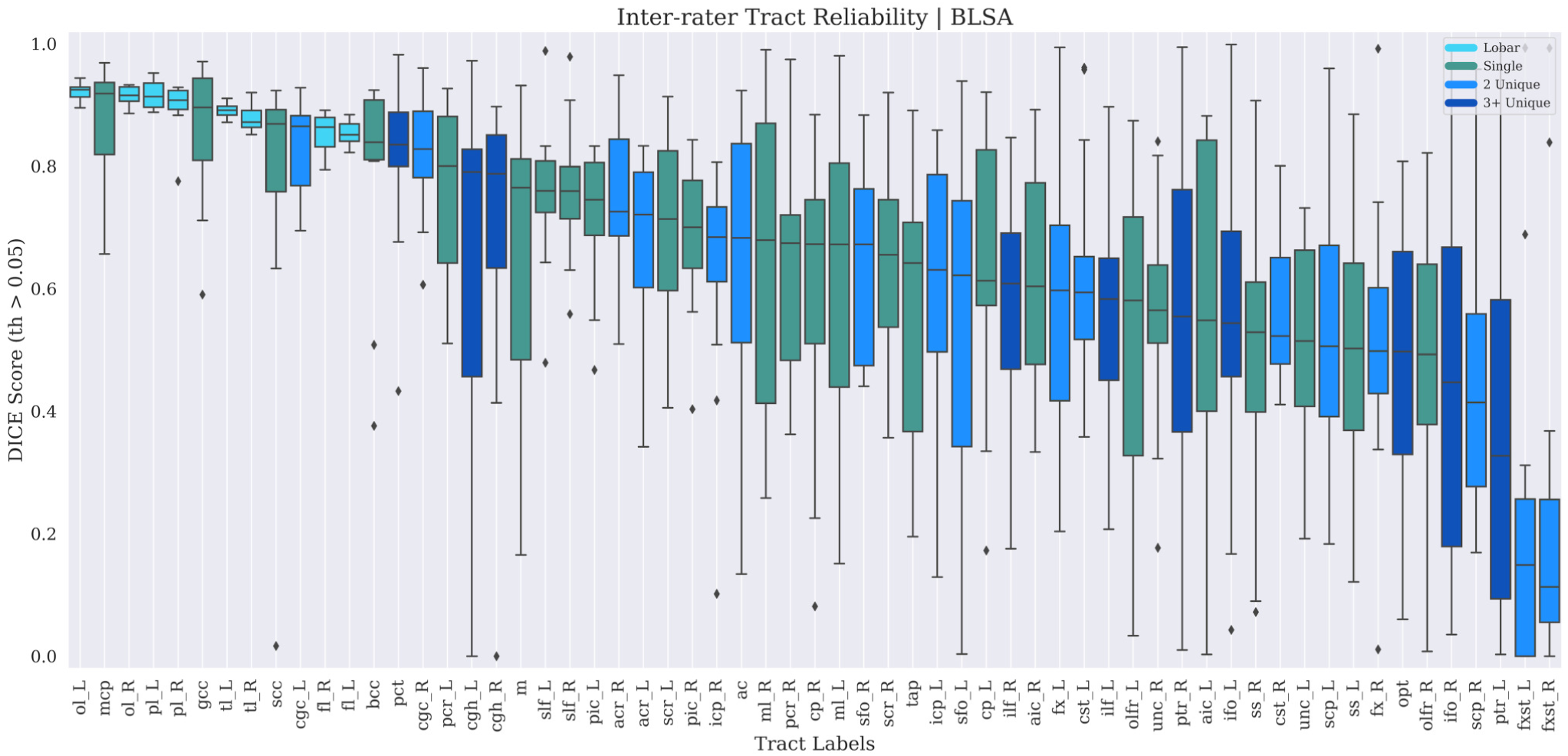
TractEM tract reliability for all 61 fiber bundles is shown. Density images were binarized with a threshold > 0.05 for similarity analysis and similarity was calculated using **DICE correlation coefficient (DICE)**. Each boxplot represents the similarity of same-subject different-rater pairs for 10 subjects from the BLSA dataset (see Section 3.3 Inter-rater Tract Reliability for the exceptions). The tracts are sorted from high to low reproducibility. Lobar regions are demonstrated with turquoise color, single label protocols are shown in dark green, protocols composed of 2-unique labels are represented by blue and protocols composed of 3+ unique labels are marked with dark blue color (see Section 2.4 TractEM Protocol Design for tract group descriptions).

**Figure B.4:**
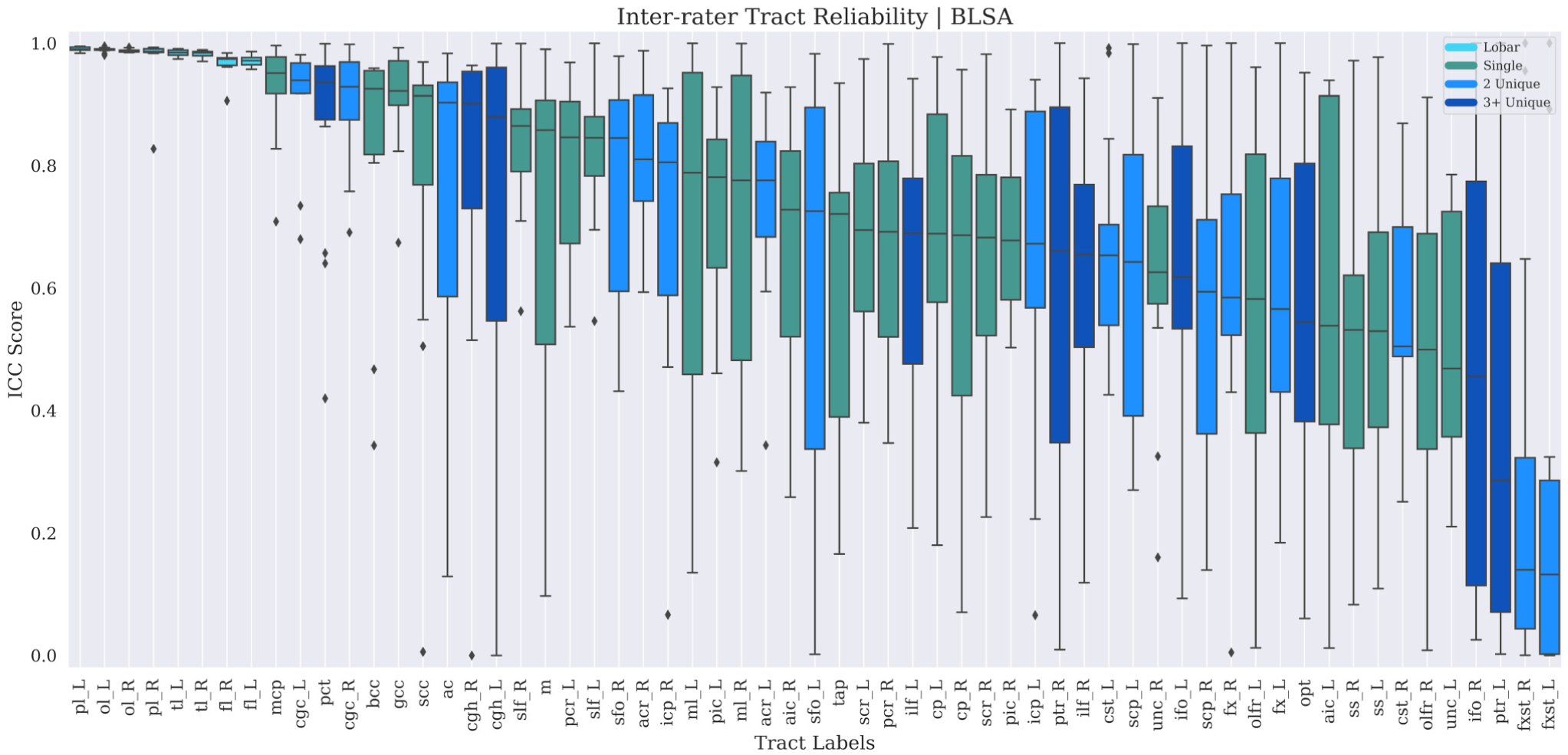
TractEM tract reliability for all 61 fiber bundles is shown. Density images were used for similarity analysis and similarity was calculated using **intraclass correlation coefficient (ICC)**. Each boxplot represents the similarity of same-subject different-rater pairs for 10 subjects from the BLSA dataset (see Section 3.3 Inter-rater Tract Reliability for the exceptions). The tracts are sorted from high to low reproducibility. Lobar regions are demonstrated with turquoise color, single label protocols are shown in dark green, protocols composed of 2-unique labels are represented by blue and protocols composed of 3+ unique labels are marked with dark blue color (see Section 2.4 TractEM Protocol Design for tract group descriptions).

**Figure B.5:**
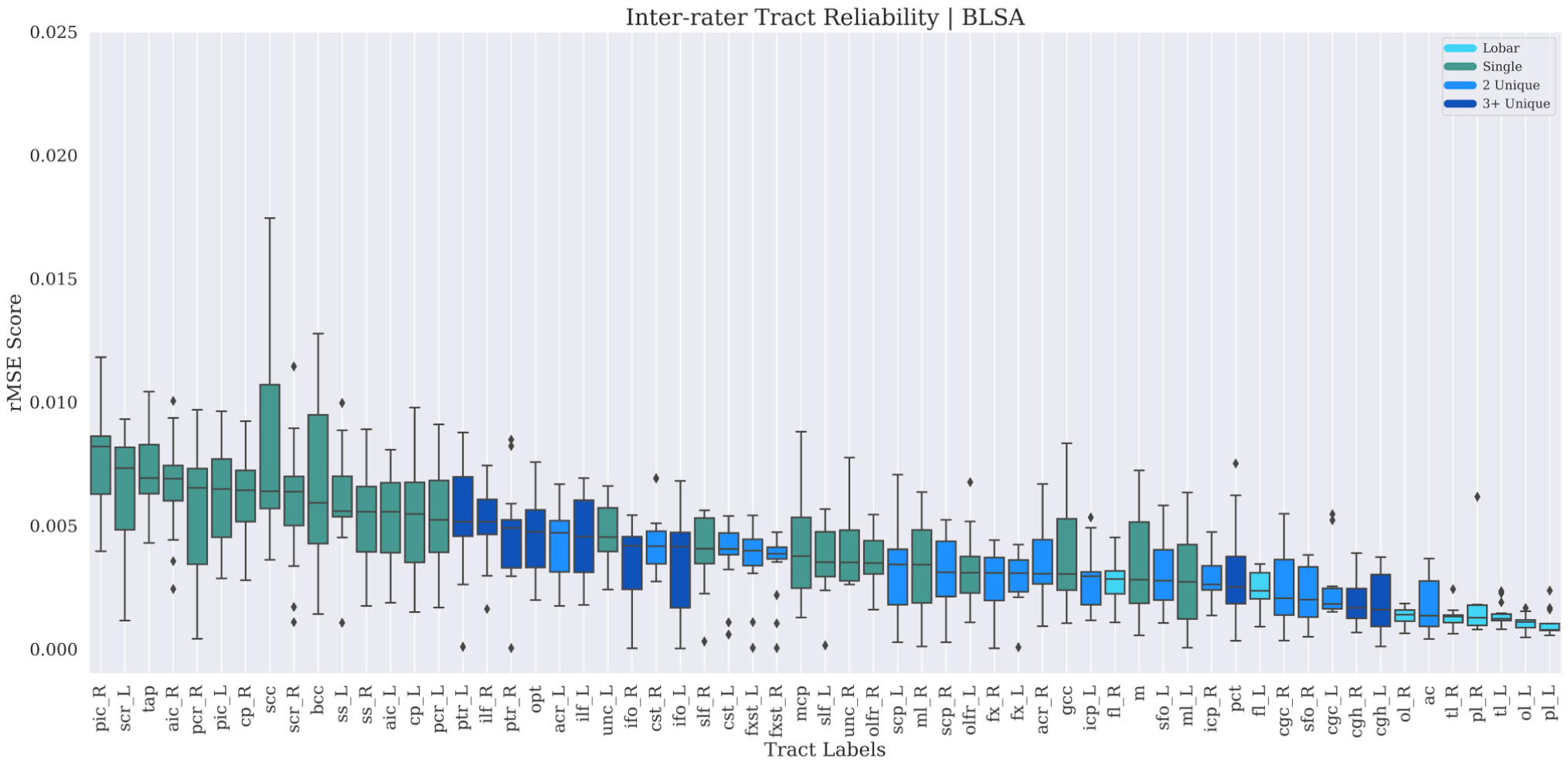
TractEM tract reliability for all 61 fiber bundles is shown. Density images used for similarity analysis and similarity was calculated using **root mean squared error (rMSE)**. Each boxplot represents the similarity of same-subject different-rater pairs for 10 subjects from the BLSA dataset (see Section 3.3 Inter-rater Tract Reliability for the exceptions). The tracts are sorted from high to low reproducibility. Lobar regions are demonstrated with turquoise color, single label protocols are shown in dark green, protocols composed of 2-unique labels are represented by blue and protocols composed of 3+ unique labels are marked with dark blue color (see Section 2.4 TractEM Protocol Design for tract group descriptions).

**Figure B.6:**
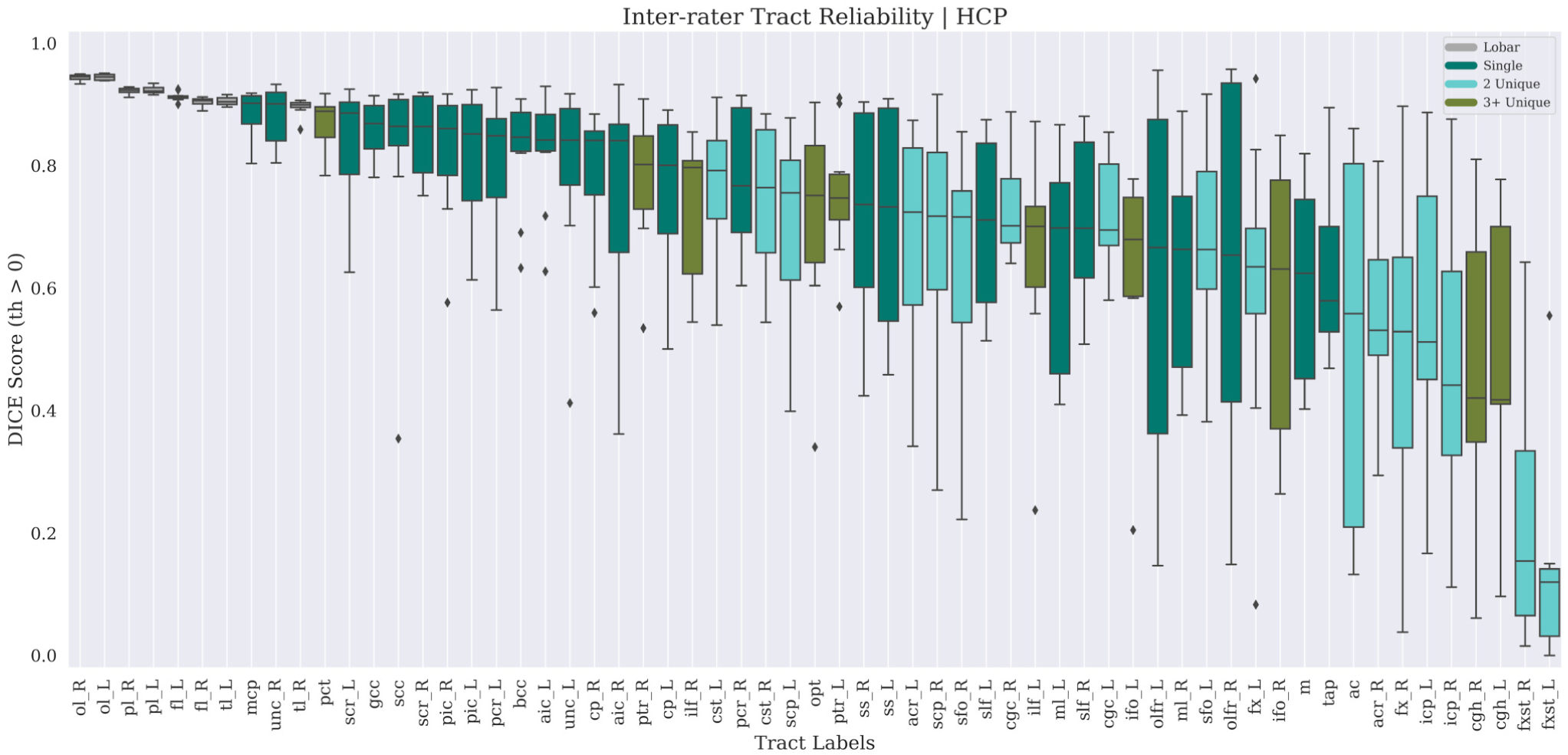
TractEM tract reliability for all 61 fiber bundles is shown. Density images were binarized with a threshold > 0 for similarity analysis and similarity was calculated using **DICE correlation coefficient (DICE)**. Each boxplot represents the similarity of same-subject different-rater pairs for 10 subjects from the HCP dataset (see Section 3.3 Inter-rater Tract Reliability for the exceptions). The tracts are sorted from high to low reproducibility. Lobar regions are demonstrated with gray color, single label protocols are shown in dark green, protocols composed of 2-unique labels are represented by cool green and protocols composed of 3+ unique labels are marked with olive color (see Section 2.4 TractEM Protocol Design for tract group descriptions).

**Figure B.7:**
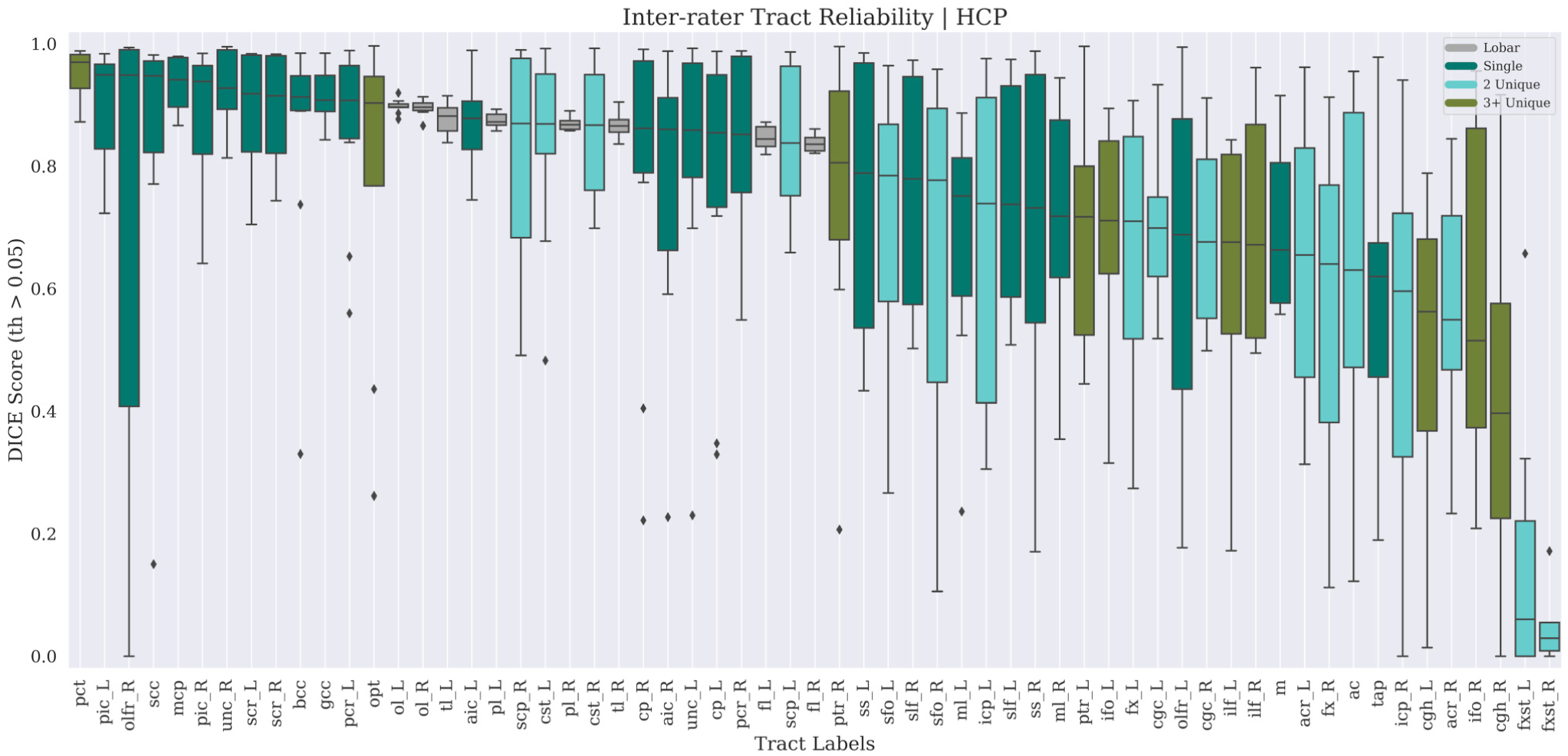
TractEM tract reliability for all 61 fiber bundles is shown. Density images were binarized with a threshold > 0.05 for similarity analysis and similarity was calculated using **DICE correlation coefficient (DICE)**. Each boxplot represents the similarity of same-subject different-rater pairs for 10 subjects from the HCP dataset (see Section 3.3 Inter-rater Tract Reliability for the exceptions). The tracts are sorted from high to low reproducibility. Lobar regions are demonstrated with gray color, single label protocols are shown in dark green, protocols composed of 2-unique labels are represented by cool green and protocols composed of 3+ unique labels are marked with olive color (see Section 2.4 TractEM Protocol Design for tract group descriptions).

**Figure B.8:**
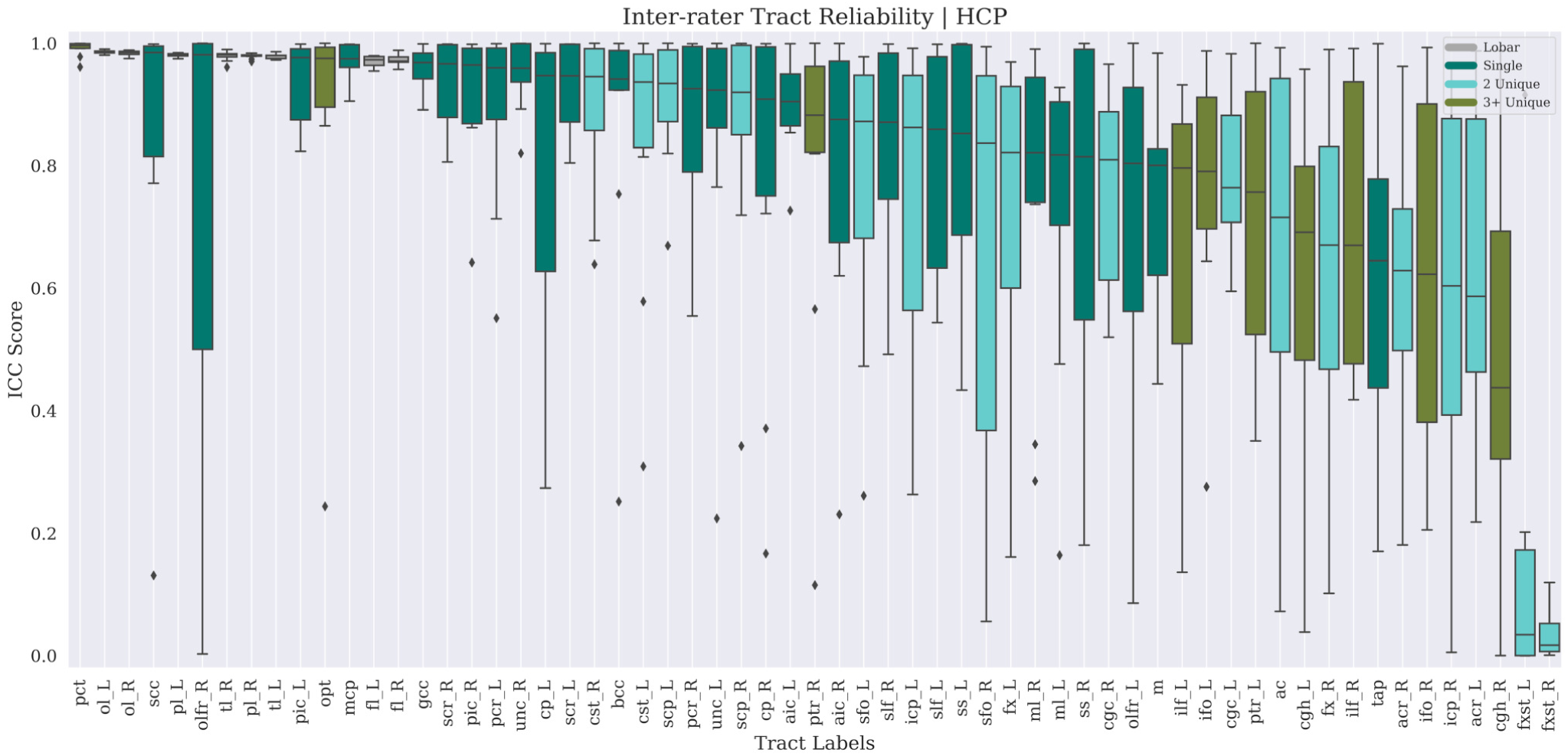
TractEM tract reliability for all 61 fiber bundles is shown. Density images were used for similarity analysis and similarity was calculated using **intraclass correlation coefficient (ICC)**. Each boxplot represents the similarity of same-subject different-rater pairs for 10 subjects from the HCP dataset (see Section 3.3 Inter-rater Tract Reliability for the exceptions). The tracts are sorted from high to low reproducibility. Lobar regions are demonstrated with gray color, single label protocols are shown in dark green, protocols composed of 2-unique labels are represented by cool green and protocols composed of 3+ unique labels are marked with olive color (see Section 2.4 TractEM Protocol Design for tract group descriptions).

**Figure B.9:**
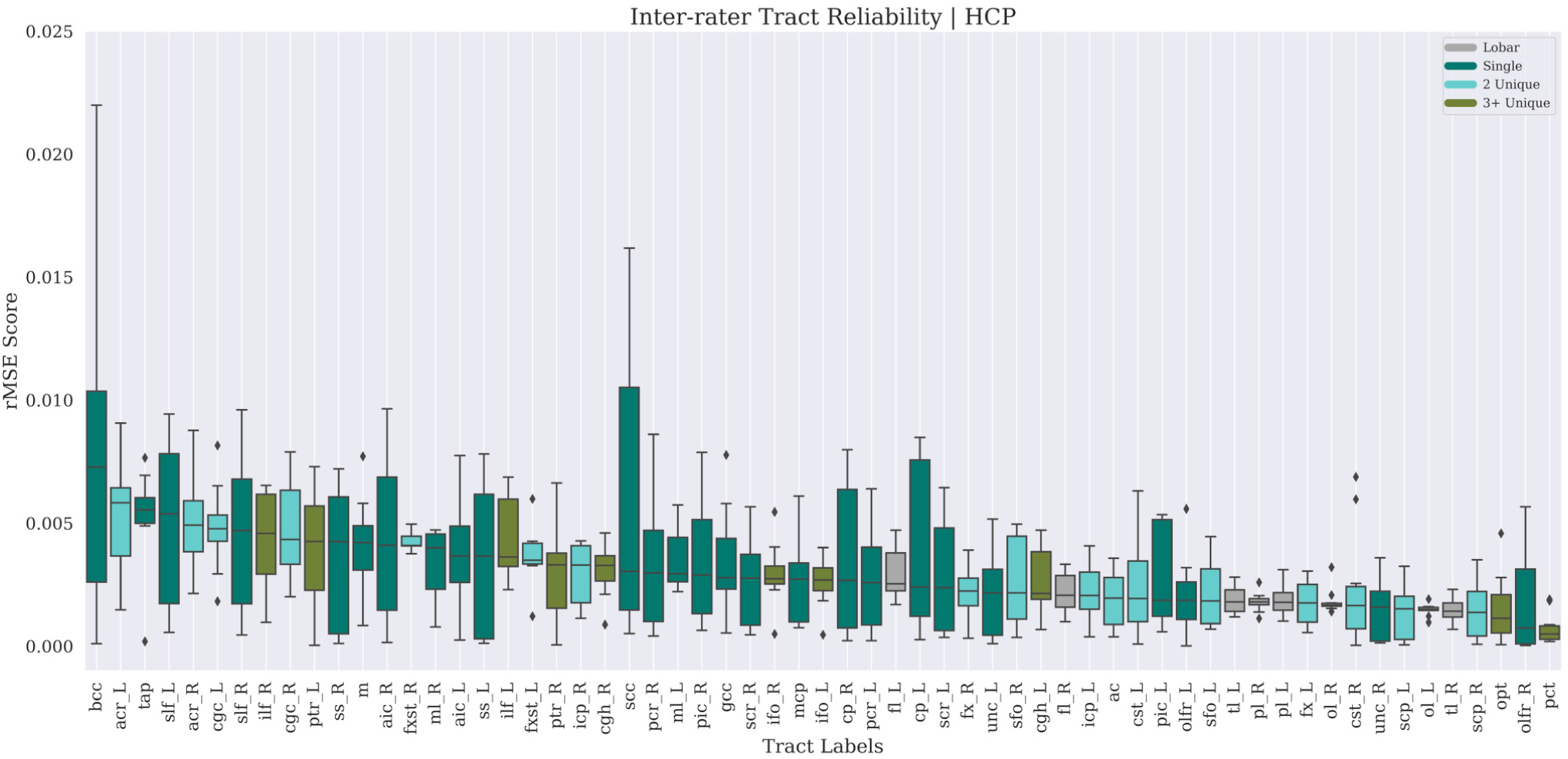
TractEM tract reliability for all 61 fiber bundles is shown. Density images were used for similarity analysis and similarity was calculated using **root mean squared error (rMSE)**. Each boxplot represents the similarity of same-subject different-rater pairs for 10 subjects from the HCP dataset (see Section 3.3 Inter-rater Tract Reliability for the exceptions). The tracts are sorted from high to low reproducibility. Lobar regions are demonstrated with gray color, single label protocols are shown in dark green, protocols composed of 2-unique labels are represented by cool green and protocols composed of 3+ unique labels are marked with olive color (see Section 2.4 TractEM Protocol Design for tract group descriptions).

**Figure B.10:**
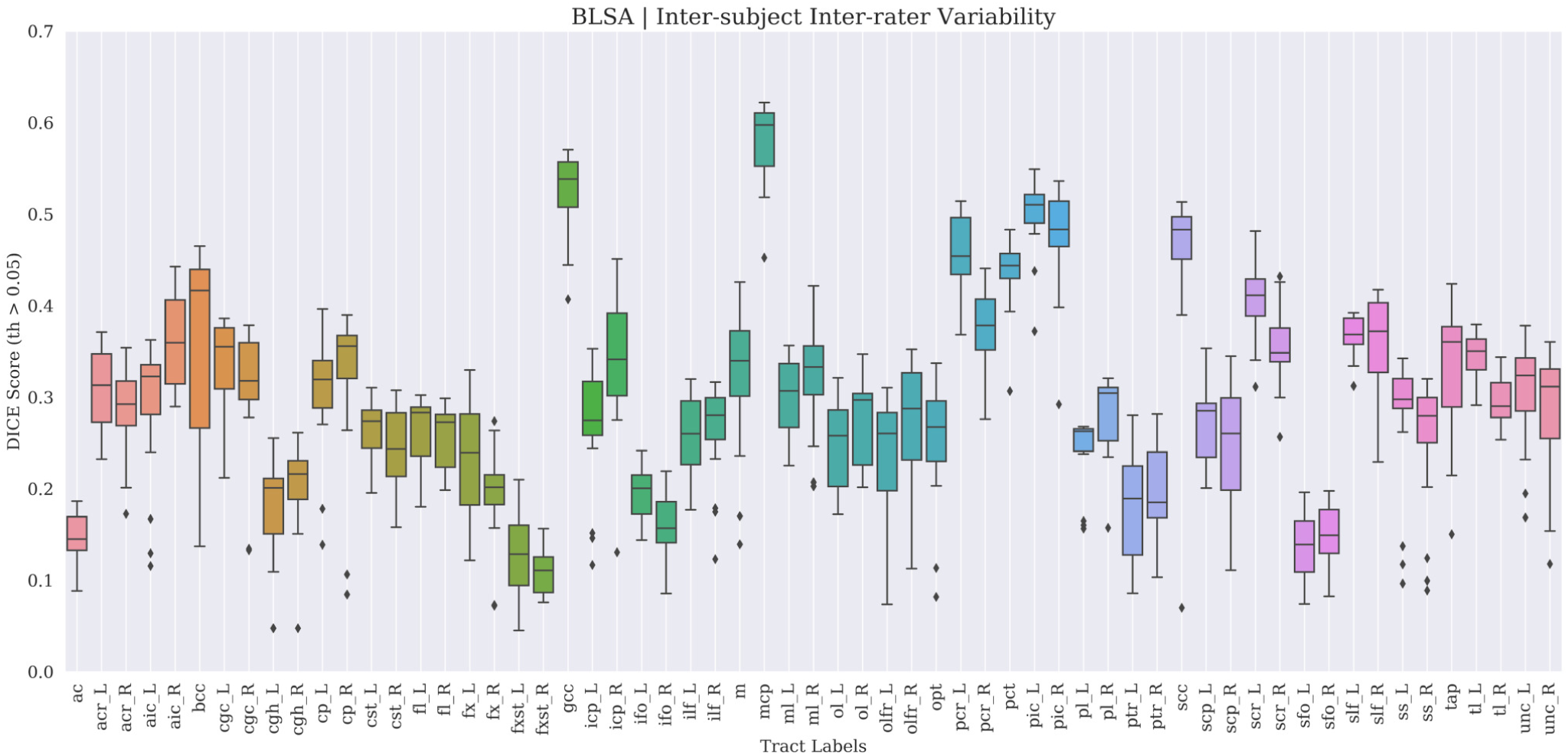
Inter-subject inter-rater variability for BLSA dataset for all tracts is shown. The reproducibility is evaluated with **DICE correlation coefficient (DICE)**. The inter-subject variability shows the anatomical variation in tracts from subject to subject. Low score implies high variation, where high score implies high correlation.

**Figure B.11:**
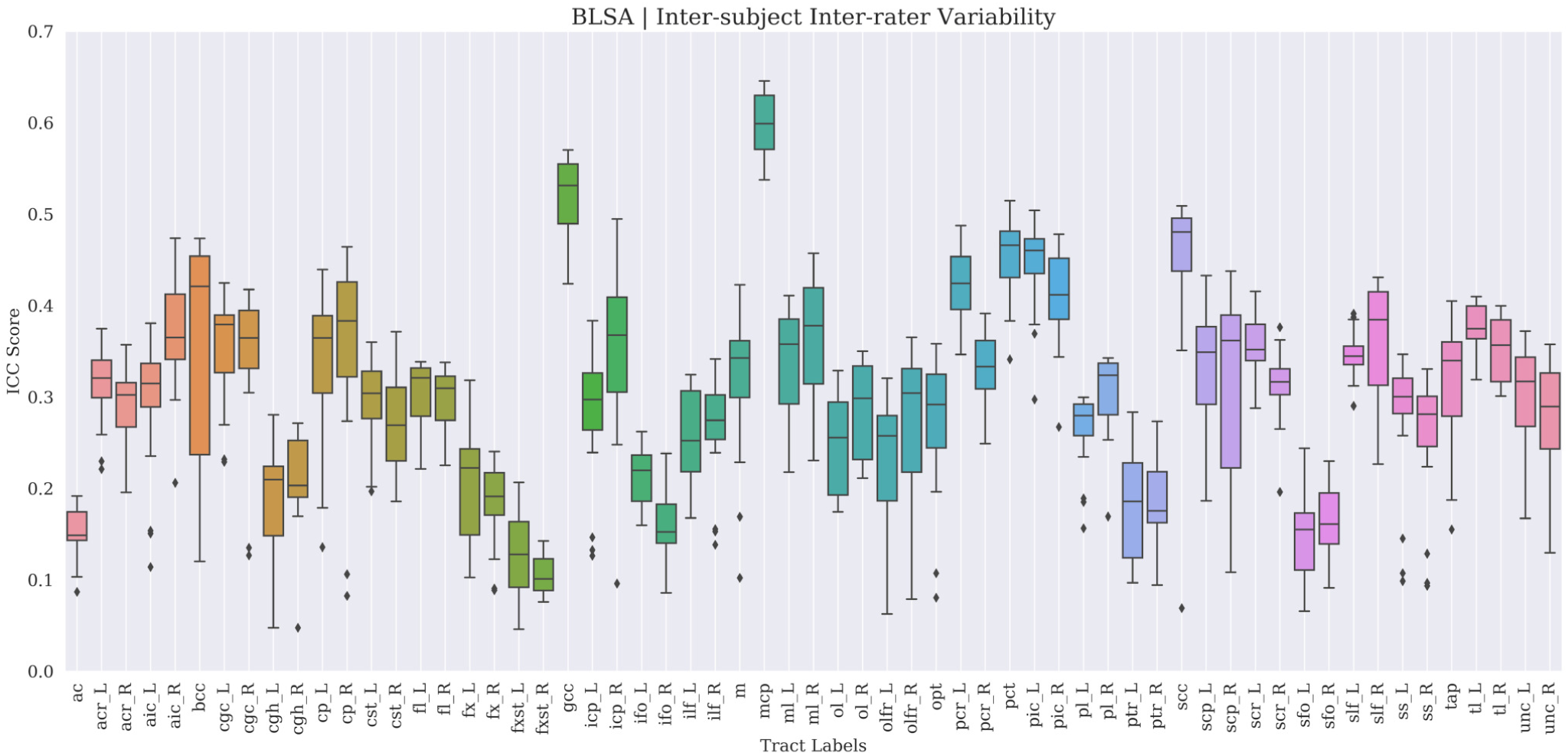
Inter-subject inter-rater variability for BLSA dataset for all tracts is shown. The reproducibility is evaluated **intra-class correlation (ICC)** measure. The inter-subject variability shows the anatomical variation in tracts from subject to subject. Low score implies high variation, where high score implies high correlation.

**Figure B.12:**
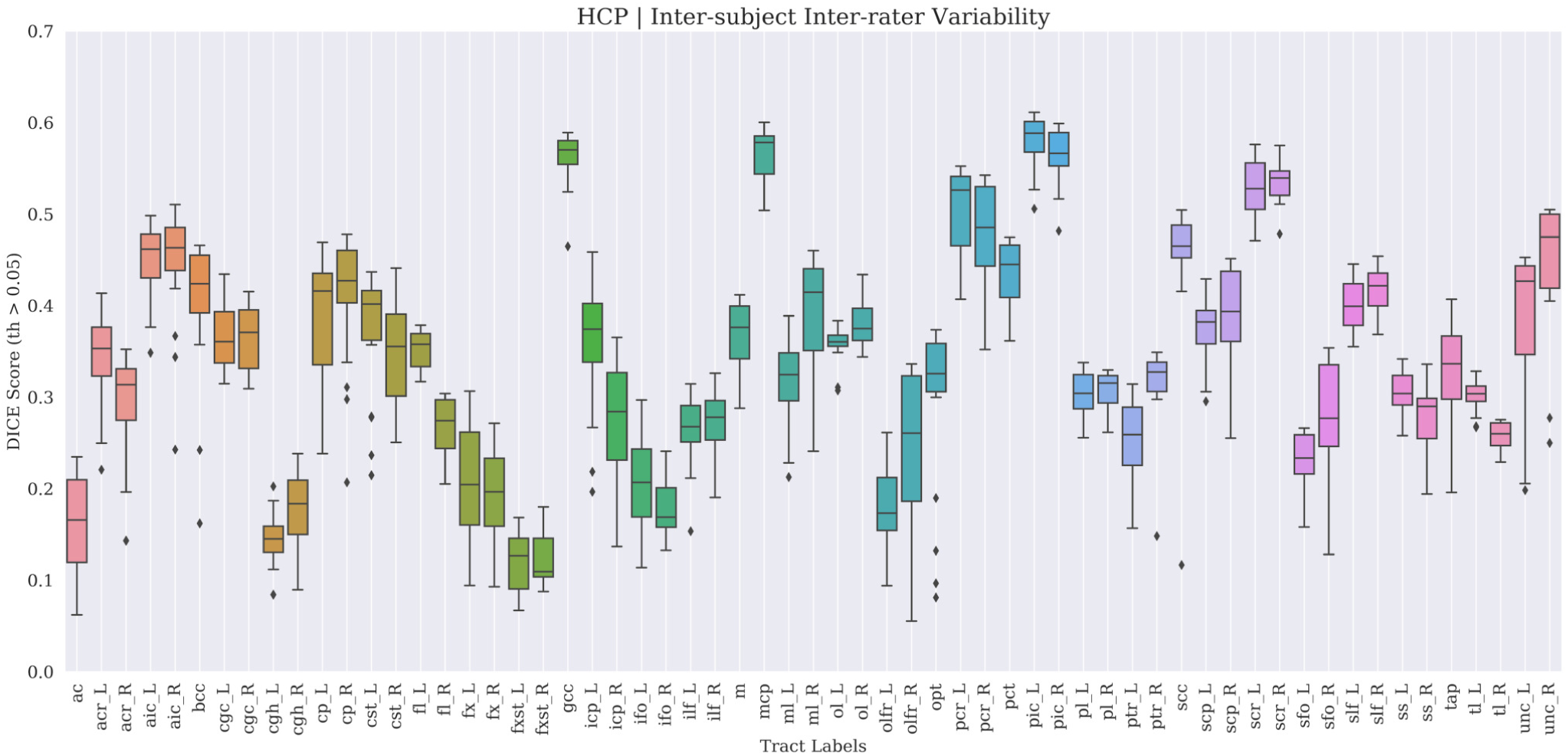
Inter-subject inter-rater variability for HCP dataset for all tracts is shown. The reproducibility is evaluated with **Dice correlation coefficient (DICE)**. The inter-subject variability shows the anatomical variation in tracts from subject to subject. Low score implies high variation, where high score implies high correlation.

**Figure B.13:**
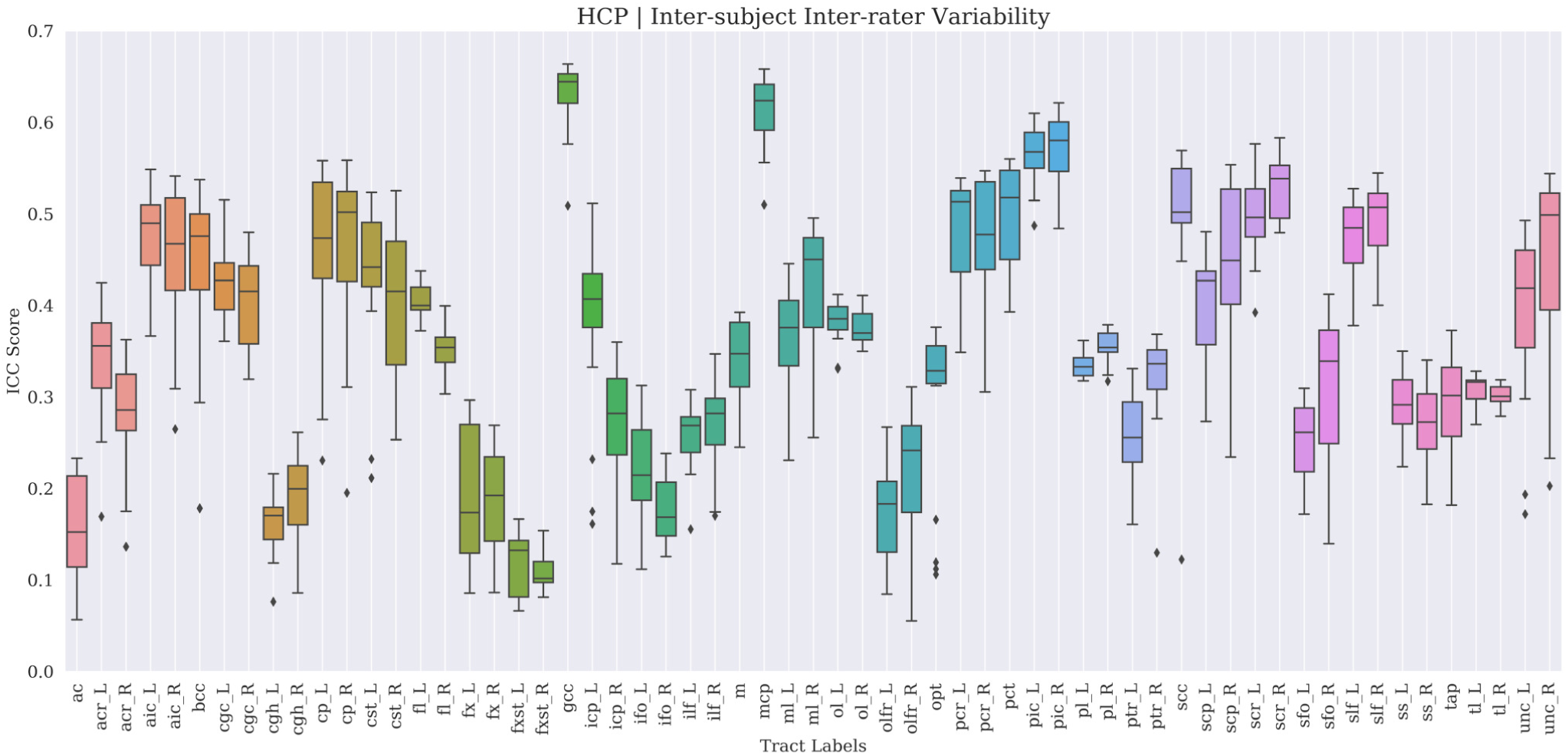
Inter-subject variability for HCP dataset for all tracts is shown. The reproducibility is evaluated with **intra-class correlation (ICC)** measure. The inter-subject variability shows the anatomical variation in tracts from subject to subject. Low score implies high variation, where high score implies high correlation.

**Figure B.14:**
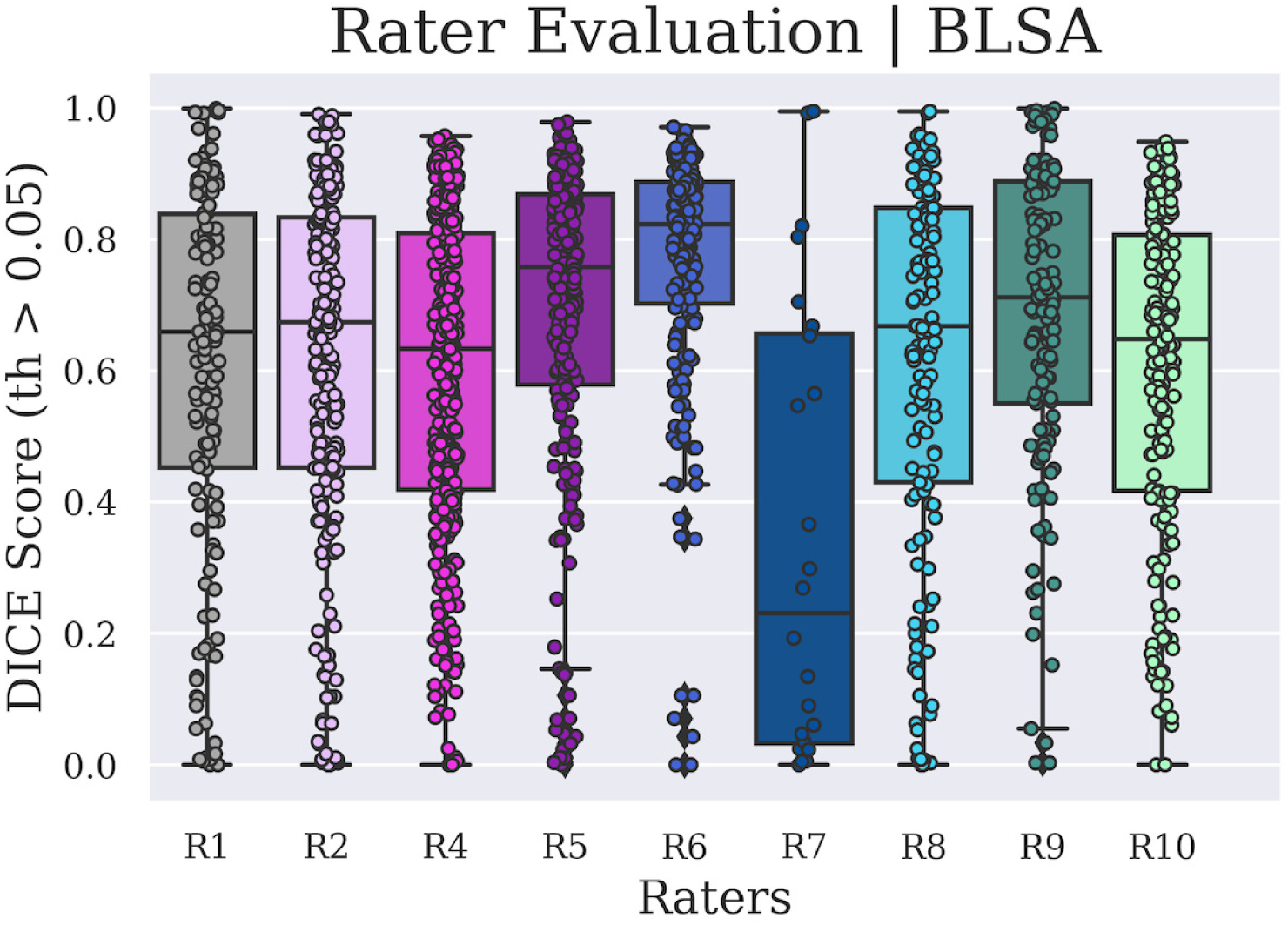
Rater evaluation for BLSA dataset is shown **DICE correlation coefficient (DICE)** with threshold > 0.05. Each data point on the boxplots indicates an intra-subject inter-rater score associated with the rater.

**Figure B.15:**
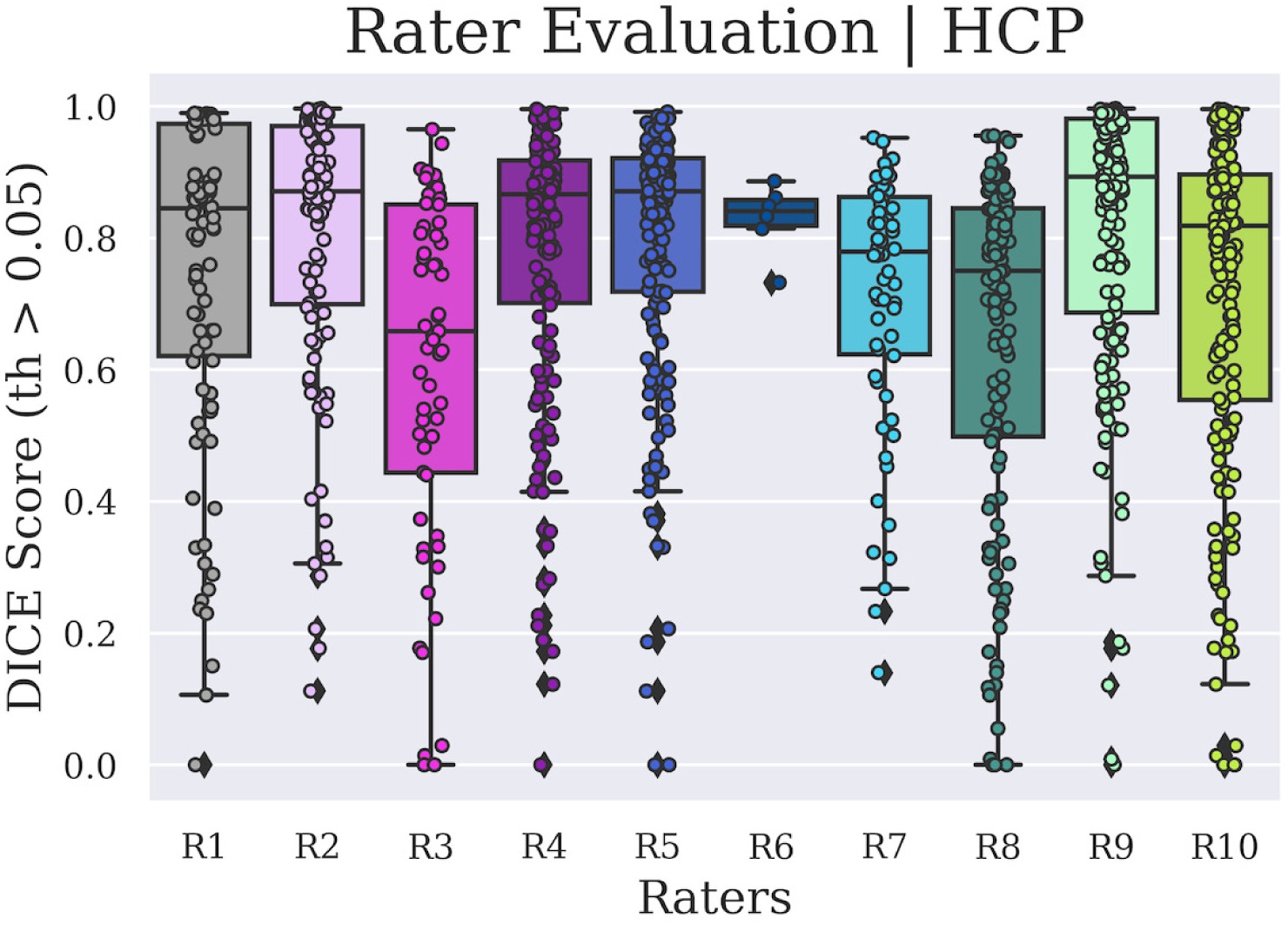
Rater evaluation for HCP dataset is shown **DICE correlation coefficient (DICE)** with threshold > 0.05. Each data point on the boxplots indicates an intra-subject inter-rater score associated with the rater.

